# Neutrophil immune profile controls spinal cord regeneration in zebrafish

**DOI:** 10.1101/2024.01.17.576035

**Authors:** Carmen de Sena-Tomás, Leonor Rebola Lameira, Patrícia Naique Taborda, Alexandre Laborde, Michael Orger, Sofia de Oliveira, Leonor Saúde

## Abstract

Spinal cord injury triggers a strong innate inflammatory response in both non-regenerative mammals and regenerative zebrafish. Neutrophils are the first immune population to be recruited to the injury site. Yet, their role in the repair process, particularly in a regenerative context, remains largely unknown. Here, we show that, following rapid recruitment to the injured spinal cord, neutrophils mostly reverse migrate throughout the zebrafish body. In addition, promoting neutrophil inflammation resolution by inhibiting Cxcr4 boosts cellular and functional regeneration. Neutrophil-specific RNA-seq analysis reveals an enhanced activation state that correlates with a transient increase in *tnf-α* expression in macrophage/microglia populations. Conversely, blocking neutrophil recruitment through Cxcr1/2 inhibition diminishes the presence of macrophage/microglia at the injury site and impairs spinal cord regeneration. Altogether, these findings provide new insights into the role of neutrophils in spinal cord regeneration, emphasizing the significant impact of their immune profile on the outcome of the repair process.

**Graphical Abstract:** 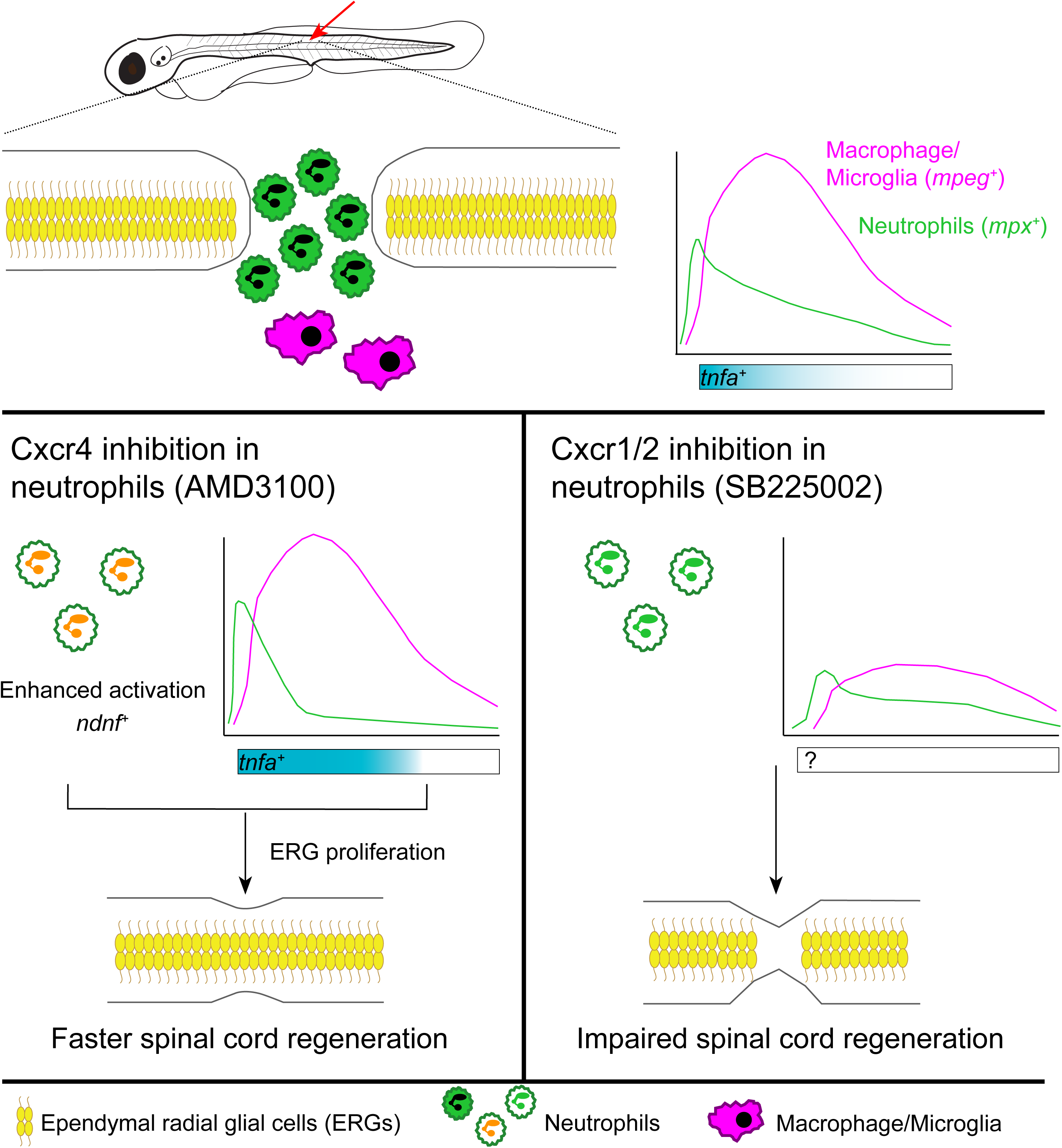

## INTRODUCTION

The immune system plays a central role in the tissue response to injury, influencing the outcome of the regenerative process in vertebrates. In mammals, spinal cord injury (SCI) elicits a chronic neuroinflammatory response that persists at the injury site [1]. This immune response starts with neutrophils, which initially support the recovery process by phagocytizing cellular debris and releasing cytokines and chemokines that promote the infiltration of macrophages and other immune cells into the damaged tissue [2, 3]. However, later on, neutrophils accumulate at the demyelinated lesion core alongside monocyte-derived macrophages, pericytes, and endothelial cells, being part of the scar that creates an inhibitory tissue microenvironment for axonal regrowth [4]. Importantly, although neutrophil infiltration into injured tissue has been widely reported, its role and contribution to SCI pathology is not well understood [5].

In contrast to mammals, zebrafish possess a remarkable regenerative capacity quickly recovering their swimming ability after SCI [6–8]. In these injured fish, the acute inflammatory response, where neutrophils are first recruited, followed by macrophages and microglia, is quickly resolved [9]. Ependymal radial glial cells (ERGs) proliferate and differentiate into new neurons and axons regrow through a repair-permissive extracellular matrix [7, 10, 11]. Importantly, pro-regenerative roles associated with the inflammatory response have been recently demonstrated [12]. These include Tnf-α secretion by macrophages to promote neurogenesis [13] and Neurotrophin-3 production by regulatory T cells to induce the proliferation of neural progenitors [14]. Besides the requirement of Il-1β, partly produced by neutrophils during the early regeneration phase, the specific role of neutrophils in spinal cord regeneration in zebrafish has not been demonstrated [9].

Neutrophils are a heterogeneous cell population, and their maturity grade and activation state significantly influence their functional outcomes. Notably, a subset of immature neutrophils has demonstrated axonal repair properties by secreting growth factors during SCI repair in mammals [15]. These transcriptionally active cells can interact with various cell types, including other leukocytes, and shape their responses accordingly. In particular, multiple mechanisms employed by neutrophils to modulate macrophage immune responses across diverse biological challenges have been described [16].

This study elucidates the impact of neutrophils on spinal cord regeneration. We demonstrate that upon quick recruitment to the injured zebrafish spinal cord, neutrophils primarily reverse migrate throughout the body to resolve their numbers at the site of injury. We further show that the modulation of neutrophil infiltration dynamics at the injury site has a profound impact on the regenerative outcome. Specifically, accelerating neutrophil inflammation resolution through Cxcr4 inhibition leads to improved regeneration of the spinal cord. Neutrophil-specific RNA sequencing (RNA-seq) analyses reveal an enhanced activation state, resulting in a transient increased expression of *tnf-α* in macrophage/microglia, along with upregulation of the *neuron-derived neurotrophic factor* (*ndnf*). Conversely, delaying neutrophil recruitment via Cxcr1/2 inhibition ablates macrophage/microglia infiltration peak and impairs the regenerative process. Overall, our study unveils the previously unknown significance of the neutrophil immune state in shaping the spinal cord microenvironment during regeneration. Gaining insight into the beneficial and detrimental roles of neutrophils at the injury site could pave the way for the development of novel therapies to treat spinal cord injuries.

## MATERIALS AND METHODS

### Zebrafish Lines and Husbandry

Zebrafish lines were raised and maintained under standard conditions [17], following FELASA recommendations [18]. The AB wild-type strain and the following transgenes were used: *Tg(mpx:GFP)^i114^* [19], *Tg(mpx:mCherry-2A-Rac2)^zf306^* [20], *Tg(mpx:mCherry-2A-Rac2D57N)^zf307^* [20], *TgBAC(mpx:GAL4.VP16)^sh267^* [21], *Tg(UAS:Kaede)^rk8^* [22] (aka *Tg(mpx:Kaede)*), *Tg(lyz:NTR-mCherry)^sh260^* [23], *Tg(mpeg:EGFP)^gl22^* [24], *Tg(mpeg1.1:mCherryF)^ump2^* [25], *TgBAC(tnfa:GFP)^pd1028^* [26], *Tg(gfap:GFP)^mi2001^* [27], *Tg(HB9:GFP)^ml2^* [28], *Tg(olig2:dsRed2)^vu19^* [29], and *Tg(fli1:EGFP)^y1^* [30].

Most experimental analyses were performed in larvae up to 5 days-old, reducing the number of animals used and adhering to the principles of the 3 Rs. All zebrafish experiments were performed in compliance with animal welfare legislation and approved by the ethical committee of the Portuguese veterinary department (license reference 0421/000/000/2021, Direcção Geral de Agricultura e Veterinária (DGAV)).

### Spinal Cord Transections and Drug Treatments

Spinal cord transections were performed in 3 days-post-fertilization (dpf) zebrafish larvae as previously described [31]. Briefly, larvae were anesthetized in 0.5mM tricaine (MS222, Sigma Aldrich), placed in the lateral position in an agarose-coated petri dish and the spinal cord was fully transected at the level of the anal pore using the tip of a 30G needle.

For drug treatments, larvae were randomly distributed into different conditions. Cxcr4 inhibition was achieved by incubating the larvae with 25 µM AMD3100 (Sigma Aldrich #239820) supplemented with 0.1% DMSO from 4 hours post-injury (hpi) until 24 hpi. For Cxcr1/2 inhibition, larvae were treated with 5 µM SB225002 (Sigma Aldrich #SML0716) plus 0.1% DMSO from one hour before spinal cord transection until 6 hpi. To induce the transition from pro-inflammatory to anti-inflammatory in *mpeg^+^* cells, larvae were incubated in 50 µM metformin in E3 media without methylene blue from 24 hpi until 48 hpi [32].

### Swimming Recovery Assay

Behavioral assessment of zebrafish larvae was performed as previously described with minor modifications [33]. 4- to 8 dpf larvae were individually placed in arenas filled with E3 without methylene blue, and their swimming activity was recorded for 90 minutes. After a 30-minute acclimatation period, three 10-minute light-dark cycles were employed to stimulate the swimming response. Experiments were automatically tracked using either a Daniovision^TM^ system with a 96-well plate arena and Ethovision X.T.8.5.614 tracking software (Noldus, Wageningen, The Netherlands) or a custom-made setup featuring circular arenas (22 mm diameter, 3.25 mm depth). In the custom setup, whole-field light and dark visual stimuli were generated using OpenGL fragment shaders and projected from below onto a flat diffusing screen at 60Hz using an Optoma ML750e LED projector. To track the fish, the set-up was illuminated from below using a custom-built lightbox with an array of 850 nm LEDs. Behavior was recorded from above the arenas using a Mikrotron EoSens®4.0MCX6-CM high-speed camera with a Schneider Xenoplan 2.0/28 objective and an 790nm colored glass long-pass filter. Images were acquired at 100 frames per second via a SiliconSoftware microEnable 5AQ8-CXP6D ironman framegrabber, and behavior data was extracted and processed online in parallel using custom written software in C# and the OpenCV library. Fish identification was achieved by subtracting a previously acquired background image from each median filter-smoothed image and using a flood fill algorithm based on the darkest pixel within each arena region. Fish location was determined as the center of mass of the blob generated by the flood fill, and position analysis was performed in MATLAB (MathWorks). Each behavior trace was smoothed out by performing a linear 2D interpolation between pairs of points more than 0.9 mm apart from each other to allow true movements to be captured without integrating small motions due to tracking noise. Finally, smoothed traces were synchronized with the visual stimulation and total travelled distance was calculated for each trial period. Total distance moved measured during the three dark periods was averaged per animal.

### Immunofluorescence, Proliferation and Apoptosis Analysis

Whole-mount immunostainings in 3 - 5 dpf larvae were performed as previously described [9] with minor modifications. Briefly, larvae were fixed in 4% paraformaldehyde/0.2% Triton X-100 overnight at 4 °C. Subsequently, they were washed with PBS plus 1% Triton X-100 (PBT), permeabilized with collagenase (2 mg/ml) for 35 minutes at 37 °C, incubated in 50mM glycine in PBT for ten minutes at room temperature, blocked in blocking solution (4% bovine serum albumin in PBT) for two hours at room temperature, and incubated with acetylated tubulin (1:300, Sigma Aldrich #T7451) in blocking solution at 4 °C overnight. Next, they were incubated with Alexa Fluor 633 goat anti-mouse IgG (1:300, Invitrogen A21050) and DAPI (1:500) for two hours at room temperature. Finally, larvae were mounted laterally in 75% glycerol prior to imaging.

Proliferation assays were performed incubating the larvae with 3mM EdU and using the Click-iT Plus EdU kit according to manufacturer’s instructions (Invitrogen, #C10640).

Neutrophil apoptosis was measured using the ApopTag Red kit (Sigma Aldrich #S7165) as previously described [34]. Briefly, larvae were fixed as described above, permeabilized with proteinase K (10 µg/ml) for 30 minutes at room temperature, washed with PBS/0.1% Tween (PBTw) prior to another 4% PFA fixation for 20 minutes at room temperature, washed again with PBTw, treated with an acetone:ethanol mix (1:2) at -20 °C for 7 minutes and washed again with PBTw. Subsequently, they were treated with the kit’s equilibration buffer for one hour at room temperature, followed by the TdT enzyme treatment for 90 minutes at 37 °C. The reaction was stopped by two 30 minutes incubations with the stop solution at 37 °C, and PBTw washes were performed prior to the antibody incubation at 4 °C overnight.

Images were acquired with a confocal laser point-scanning Zeiss LSM 880 microscope using a Plan-Apochromat 0.8/20x air and a LD C-apochromat Crr 1.10/40x water objectives. Z-stacks were analyzed using IMARIS 9.5 (Oxford Instruments) and ImageJ (version 1.53k, NIH).

### Neutrophil Photoconversion and Live Imaging

Photoconversion of neutrophils at the injury site was performed on 3 dpf *TgBAC(mpx:GAL4.VP16)^sh267^*; *Tg(UAS:Kaede)^rk8^*; *Tg(gfap:GFP)^mi2001^* anesthetized larvae embedded in 1% low melting point agarose in E3 to maintain the proper position. In addition, E3 was supplemented with 0.16mg/ml tricaine to maintain a moisturized environment during the acquisition. For the photoconversion, larvae were placed in a Zeiss LSM880 confocal microscope under a LD C-Apochromat 40x objective. A 405-nm laser was focused into a rectangular area restricted to the injury in the spinal cord with 40% transmission, 50 iterations and 1.02 µs pixel dwell time. Successful photoconversion was assessed by the loss of fluorescence emission following excitation at 488 nm and the gain of fluorescence emission following 561 nm excitation. Thirty minutes after photoconversion, neutrophil movement was tracked in a Zeiss Cell Observer Spinning Disk confocal microscope equipped with a Evolve 512 EMCCD camera, using an EC Plan-Neofluar Ph1 0.3/10x air objective. Whole-larvae time-lapse movies were assembled from 6 x 1 tiles with 5 µm optical sections z-stacks, acquired every two minutes at a 512 x 512 resolution for six hours.

### Quantification and Statistical Analysis

All quantifications were performed blinded. Unless otherwise indicated, experiments were repeated at least three times. Lateral view maximal projections are always shown rostrally to the left and caudally to the right. Neutrophils, macrophage/microglia and *tnf-α^+^* cells were quantified using IMARIS 9.5 (Oxford Instruments). A fixed rectangular region of interest (ROI) was adjusted for each image: 650 µm x 55 µm to quantify the cells in the caudal hematopoietic tissue, 250 µm x 170 µm to quantify the photoconverted neutrophils at the injury site, and an injury-dependent area restricted to the spinal cord including 50 µm rostrally and caudally from the border. Within this region, cells were detected applying an intensity threshold cutoff, followed by a surface filtering. EdU incorporation quantifications were performed as previously described [7] by manually analyzing all consecutive images in z-stacks within a 210 µm length containing the injury site. Neutrophil-specific apoptosis was assessed by manually quantifying the GFP^+^ neutrophils labelled with TUNEL 100 µm rostrally and caudally from the border. Axonal bridge was considered if there was a continuity of labelling between the two spinal cord stumps and its thickness was measured using ImageJ (NIH). Percentage reduction in neutrophil numbers at the injury site were calculated as previously described [35]. Statistical analysis was performed using GraphPad Prism 8. One-way ANOVA followed by Tukey’s or Dunnet’s multiple comparison post-test, two-way ANOVA followed by Tukey’s multiple comparison post-test and unpaired two-tailed t-test were used, as indicated in the figure legends. The following nomenclature was employed to present results: ns, not significant, *p < 0.05, **p < 0.01; *** p < 0.001, **** p < 0.0001. Statistical values in bar graphs are displayed as mean ± standard error of the mean (SEM), while in violin plots median and quartiles are marked. Figures were prepared with Adobe Photoshop CS6, Adobe Illustrator CS6 and Adobe InDesign CS6, and graphs were generated using GraphPad Prism 8.

### Tissue Dissociation and FACS

Trunks containing the lesion site were dissociated following an unpublished protocol from Yi Feng’s lab. Briefly, anesthetized larvae were dissected and placed in dissociation solution (2.5 mg/ml collagenase IV, 15 mM HEPES, 25 mM D-Glucose, 2% goat serum in HBSS without magnesium and calcium), incubated at 28.5 °C shaking at 300rpm, and vigorously dissociated using a pipette. Subsequently, they were filtered through a 40 µm strainer (BD Biosciences) and centrifuged at 300 g for 10 minutes at 4 °C. Next, cells were resuspended in incubation media (dissociation solution without collagenase IV) and stained for viability using DAPI (1µl/ml) for 10 min at 4 °C. Single-cell suspensions obtained from 100 trunks (RNA-seq) or 15 trunks (qPCR) were used to sort GFP^+^ cells directly into TRIzol using a FACSAria III (BS Biosciences). All samples were processed in less than two hours.

### Quantitative PCR and RNA Sequencing

For qPCR, total RNA was extracted from thirty pooled trunks containing the injury site for each condition and timepoint analyzed using TRIzol reagent (Invitrogen) and RNeasy Mini Kit (Qiagen #74104). The purified RNAs were reverse transcribed with the iScript^TM^ Reverse Transcription Supermix (Bio-Rad #1708841). qPCR was performed with Power SYBR Green PCR Master mix (Applied Biosystems #4368702) and a 7500 Fast Real-time PCR system (Applied Biosystems). All experiments were completed using at least three biological replicates, and expression levels were normalized to *gapdh* or *actb1*. Primer sequences are noted in Supplemental Table 1. Bulk RNA Sequencing was performed at Single Cell Discoveries (Utrecht, The Netherlands) using an adapted version of the CEL-seq protocol. Briefly, total RNA was extracted from 2500 – 4800 GFP^+^ cells using the standard TRIzol (Invitrogen) protocol and used for library preparation and sequencing. mRNA was processed as described previously, following an adapted version of the single-cell mRNA seq protocol of CEL-Seq [36, 37]. Samples were barcoded with CEL-seq primers during the reverse transcription and pooled after second strand synthesis. The resulting cDNA was amplified with an overnight *in vitro* transcription reaction and the quality of the RNA was evaluated using TapeStation (Agilent). From this amplified RNA, sequencing libraries were prepared with TruSeq small RNA primers (Illumina) and paired-end sequenced on a Nextseq™ 500 (Illumina), high output, with a 1x75 bp Illumina kit (Read 1: 26 cycles, index read: 6 cycles, Read 2: 60 cycles).

### RNA-sequencing Data Processing and Analysis

Read 1 was used to identify the Illumina library index and CEL-Seq sample barcode and read 2 was aligned to the GRCz11 reference transcriptome using Burrow-Wheeler aligner (BWA-MEM) [38]. Reads that mapped equally well to multiple locations were discarded. Mapping and generation of count tables was done using the MapAndGo script [39]. Normalization and differential gene expression analyses were conducted using DESeq2 v1.38 [40]. For PCA visualization, the data was transformed using regularized log transformation to ensure an equal contribution from all genes. Log2 fold changes were adjusted using Bayesian shrinkage [41]. Significant differentially expressed genes between conditions were selected using a p-adjusted value < 0.05 in R (version 4.2.2). Venn overlaps of significantly DEGs were generated with the VennDiagram package [42]. Functional enrichment was performed using g:Profiler [43], employing default parameters. KEGG pathways plot was created using the gplots package (version 3.1.3). Network reconstructions were performed by retrieving protein-protein interacting relationships using STRING (version 11.5, default parameters, https://string-db.org). For networks associated with KEGG pathways, interactions only between genes associated with specified KEGG pathways were examined.

### Data and Software Availability

The RNA sequencing data reported in this paper is available under accession number XXXXX at the Gene Expression Omnibus. All other relevant data may be requested from the corresponding authors.

## RESULTS

### Neutrophils recruited to the injured spinal cord reverse migrate

Neutrophils were previously described as the first immune population to infiltrate the spinal cord after injury [9]. However, their recruitment and clearance dynamics are influenced by the injury severity [44]. To characterize these processes in our zebrafish larvae injury model, we took advantage of the Tg(*mpx:GFP*) line, which specifically labels neutrophils [19], to quantify their presence at the injury site at different timepoints (Figure 1A–D). Following complete transection of the spinal cord of 3-days post-fertilization (3 dpf) larvae, a rapid neutrophil recruitment was observed, which peaked between 4- and 6-hours post-injury (hpi) (24 ± 6 and 26 ± 7, respectively). Subsequently, neutrophil infiltration was resolved, and the numbers at the injury site returned to uninjured levels by 72 hpi (3 ± 2) (Figure 1E). In correlation with this infiltration peak, a 76% reduction in the number of neutrophils in the caudal hematopoietic tissue (CHT) was detected at 4 hpi, which returned to uninjured levels by 24 hpi (Figure 1F). This aligns with the fact that the CHT is the main reservoir of neutrophils in zebrafish larvae and it is in close proximity to the injury site [45].

**Figure 1:**
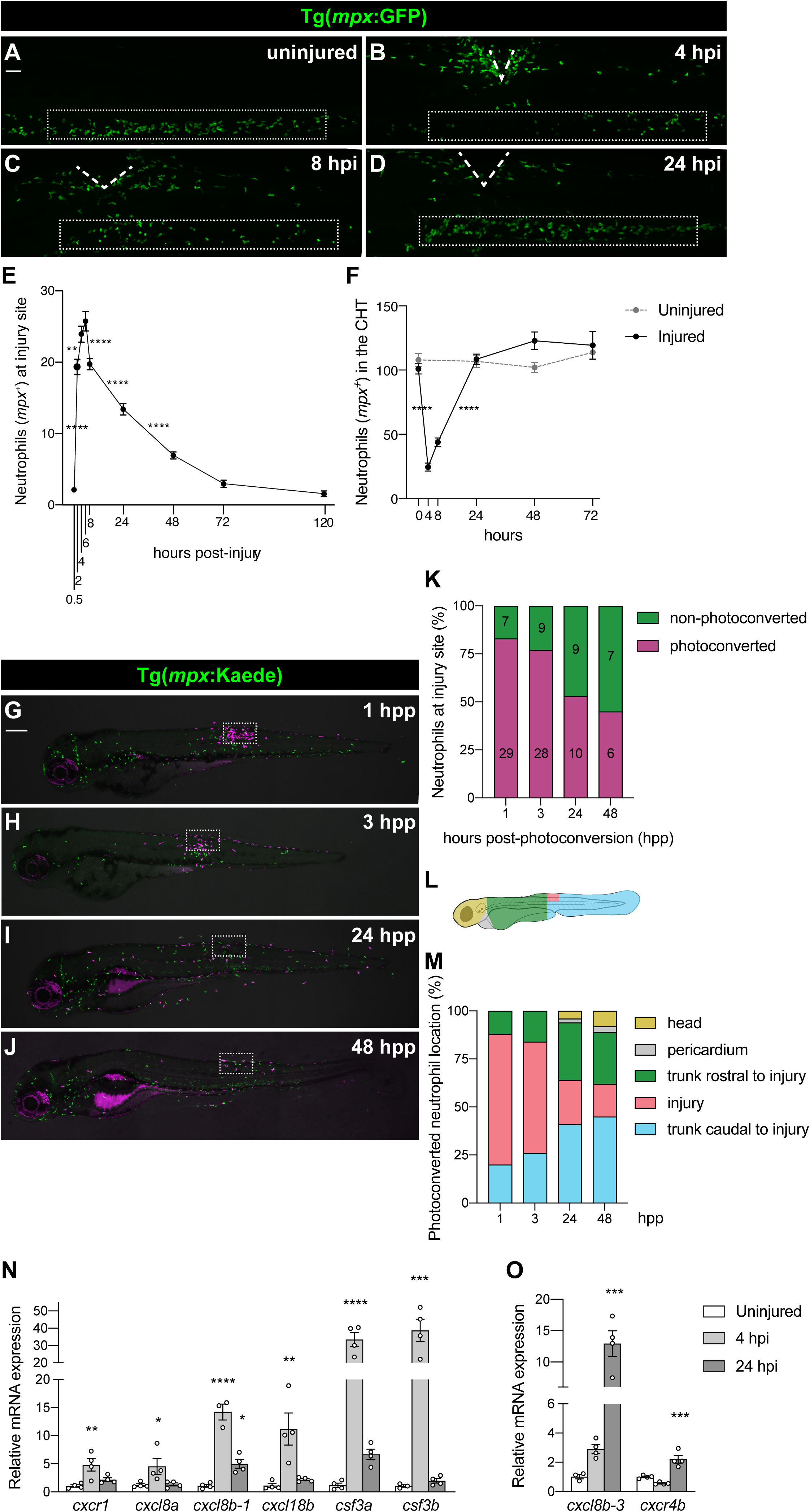
Neutrophils recruited to the injured spinal cord through conserved signaling pathways reverse migrate. **(A - D)** Representative lateral images of 3 dpf uninjured and injured larvae highlighting the recruitment, accumulation and gradual departure of neutrophils (*mpx^+^*). Bellow the injury site, the caudal hematopoietic tissue (CHT) initially empties and the replenishes, mirroring the neutrophil movements in the spinal cord. The injury site is marked with dashed lines and the CHT is outlined in the rectangular boxes. Scale bar, 50 μm. **(E)** Quantification reveal the number of *mpx^+^* cells at the injury site, peaking at 4 – 6 hours post-injury (hpi). Each square represents mean ± SEM (n = 20 - 31 performed as two independent experiments). One-way ANOVA followed by Tukey’s multiple comparisons test shows a significant difference between 0.5 and 2 (p < 0.0001), 2 and 4 (p = 0.0061), 6 and 8 (p < 0.0001), 8 and 24 (p < 0.0001), and 24 and 48 hpi (p < 0.0001) with no significant difference between 4 and 6 (p = 0.8642), 48 and 72 (p = 0.0885), and 72 and 120 hpi (p = 0.9895). **(F)** Quantification of *mpx^+^* cells at the CHT demonstrated a sharp decrease between 0 – 4 hpi followed by recovery at 24 hpi. Each square represents mean ± SEM (n = 15 - 23 performed as two independent experiments). One-way ANOVA followed by Tukey’s multiple comparison test shows a significant difference between 0 and 4 (p < 0.0001) and 8 and 24 hpi (p < 0.0001) with no significant difference between 4 and 8 (p = 0.2610), 24 and 48 (p = 0.5644), 48 and 72 hpi (p > 0.9999), 0 hpi and 0 hours uninjured (hu) (p = 0.9929), 24 hpi and 24 hu (p > 0.9999), 48 hpi and 48 hu (p = 0.1294), and 72 hpi and 72 hu (p = 0.9996). **(G – J)** Representative maximal projections of injured 3 dpf larvae with neutrophils (magenta) photoconverted at the injury site at 4 hpi and then tracked live at 1, 3, 24 and 48 hours post-photoconversion (hpp). The injury site is outlined in rectangular boxes. Scale bar, 200 μm. **(K)** Quantification demonstrates the change in the ratio of photoconverted and non-photoconverted neutrophils at the injury site at 1, 3, 24 and 48 hpp (n = 13 - 15, performed as three independent experiments). Mean neutrophil numbers are displayed within the graph. **(L)** Scheme illustrating the different areas quantified to analyze the migratory pattern of the photoconverted neutrophils after spinal cord injury. **(M)** Quantification reveals the location of the photoconverted neutrophils at 1, 3, 24 and 48 hpp (n = 13 - 15, performed as three independent experiments). **(N)** Relative gene expression, as measured by qPCR, shows the upregulation of chemokines involved in guiding neutrophil forward migration. Each dot represents a biological replicate obtained from a pool of thirty trunks and each bar depicts the mean ± SEM. One-way ANOVA followed by Dunnet’s multiple comparisons test reveals significant differences between uninjured and 4 hpi samples for *cxcr1* (p = 0.0065), *cxcl8a* (p = 0.0379), *cxcl8b-1* (p < 0.0001), *cxcl18b* (p = 0.0038), *csf3a* (p < 0.0001) and *csf3b* (p = 0.0005), and between uninjured and 24 hpi for *cxcl8b-1* (p = 0.015). **(O)** Relative gene expression, as measured by qPCR, indicates changes in signaling molecules involved in neutrophil inflammation resolution. Each dot represents a biological replicate, and each bar depicts the mean ± SEM. One-way ANOVA followed by Dunnet’s multiple comparisons test shows significant differences between uninjured and 24 hpi samples for *cxcl8b-3* (p = 0.0005) and *cxcr4b* (p = 0.0008).

Next, we analyzed neutrophil behavior during the resolution phase following a spinal cord injury. Different injury models performed in the fin and heart of 3 dpf larvae have shown that neutrophils primarily undergo reverse migration to resolve their numbers at lesion sites [45–47]. Therefore, we employed the double transgenic zebrafish line *TgBAC(mpx:GAL4.VP16)*; *Tg(UAS:Kaede)* (aka Tg(*mpx:Kaede*)) [21, 22], in which neutrophils express the photoconvertible protein Kaede. Neutrophils recruited to the injury site were photoconverted at 4 hpi, coinciding with the infiltration peak, and then neutrophil behavior was assessed by non-invasive time-lapse confocal microscopy imaging for six hours (Supplemental Movie 1). As expected, we observed neutrophils reverse migrating from the injury site to different locations within the larvae. To analyze this behavior over a longer period, we repeated the photoconversion experiment and acquired images at different timepoints starting at 1-hour post-photoconversion (hpp), when neutrophil recruitment was still at its peak, until 48 hpp (Figure 1G–J). First, we analyzed the abundance of photoconverted neutrophils at the injury site, which decreased from 29 ± 9 (representing 83 ± 7%) at 1 hpp to 6 ± 2 (corresponding to 45 ± 9%) at 48 hpp. Interestingly, these quantifications reveal that neutrophils that had been photoconverted at the injury site at 4 hpi were still detected in the wound 48 hours later (Figure 1K). Next, to understand the migratory pattern of the photoconverted neutrophils and evaluate the preference of neutrophil relocation following reverse migration, we divided the zebrafish larvae into five different anatomical regions and quantified the neutrophils (Figure 1L). At 1 and 3 hpp, more than half of the photoconverted neutrophils were still present at the injury site (68 ± 12% and 58 ± 15%, respectively), and most of the reversed migrated neutrophils were found in the trunk, accounting for 11 ± 4% and 16 ± 8% rostrally to the injury and 20 ±9 % and 26 ± 9% caudally to the injury, respectively (Figure 1M). As expected, by 24 hpp, the number of photoconverted neutrophils present at the injury site decreased to 23 ± 9%. In addition, neutrophils that had migrated away from the injury could be found in more distant locations such as the head (4 ± 2%) and pericardium (1 ± 1%), although more than two-thirds were detected in the trunk (30 ± 6% and 41 ± 9% representing rostral and caudal regions, respectively) (Figure 1M). Finally, by 48 hpp, the number of photoconverted neutrophils that were still detected at the injury was reduced to a 17 ± 4%, and an increase in the number of reverse migrated neutrophils in the head (8 ± 4%) and the pericardium (3 ± 3%), while no major alterations in trunk numbers were observed (27 ± 10% and 45 ± 9% indicating rostral and caudal values respectively) (Figure 1M). Taken together, these results indicate that following spinal cord injury, neutrophils reverse migrate throughout the zebrafish larvae, reaching more distant locations as neutrophil resolution evolves, but without a clear organ preference.

Chemokine gradients and signaling guide neutrophil infiltration into damaged tissues [48]. Zebrafish Cxcr1 and Cxcr2 play redundant roles during neutrophil recruitment, Cxcr4 participates in their retention at the wound, and Cxcr2 is required for the resolution of neutrophil inflammation [35, 49, 50]. Therefore, to validate whether conserved signaling pathways control neutrophil dynamics in our spinal cord injury model, we isolated tissue containing the injury site at 4 and 24 hpi and performed qPCR. As expected, at 4 hpi and correlated with the neutrophil peak at the injury site, we observed the upregulation of *cxcr1* and its ligand *cxcl8a* [50], *cxcl8b-1* and *cxcl18b* (Cxcr2 ligands) [51–53], and *csf3a* and *csf3b* (Csf3r ligands) [54], all of which are known to be involved in neutrophil recruitment (Figure 1N). Moreover, at 24 hpi, when neutrophil infiltration was being resolved, *cxcl8b-3* (Cxcr2 ligand) and *cxcr4b* were found to be upregulated (Figure 1O). This indicates that conserved chemokine pathways control neutrophil infiltration dynamics in a spinal cord injury context.

### Delaying neutrophil recruitment to the injured spinal cord impairs regeneration

Given that our data demonstrates upregulation of chemokines such as *cxcl8a, cxcl8b-1*, and *cxcl18b* in the injured spinal cord and that these are sensed by the Cxcr1/2 receptors controlling neutrophil forward migration, we attempted to alter neutrophil recruitment using the well-characterized Cxcr1/2 antagonist, SB225002 [45, 52, 55]. SB225002 was administered one-hour prior to injury until 6 hpi to modulate the neutrophil infiltration phase (Figure 2A). As a result, we observed a 2-hour delay in neutrophil recruitment and a significant decrease in neutrophil numbers at the initial timepoints compared to DMSO-treated larvae (Figure 2B-J). Moreover, while control larvae exhibited a typical fast resolution of neutrophil infiltration, SB225002-treated larvae showed lower neutrophil resolution with no significant decrease in neutrophil count (Figure 2J, K). This indicates that our treatment not only affects neutrophil recruitment but also neutrophil behavior during the resolution process [49].

**Figure 2:**
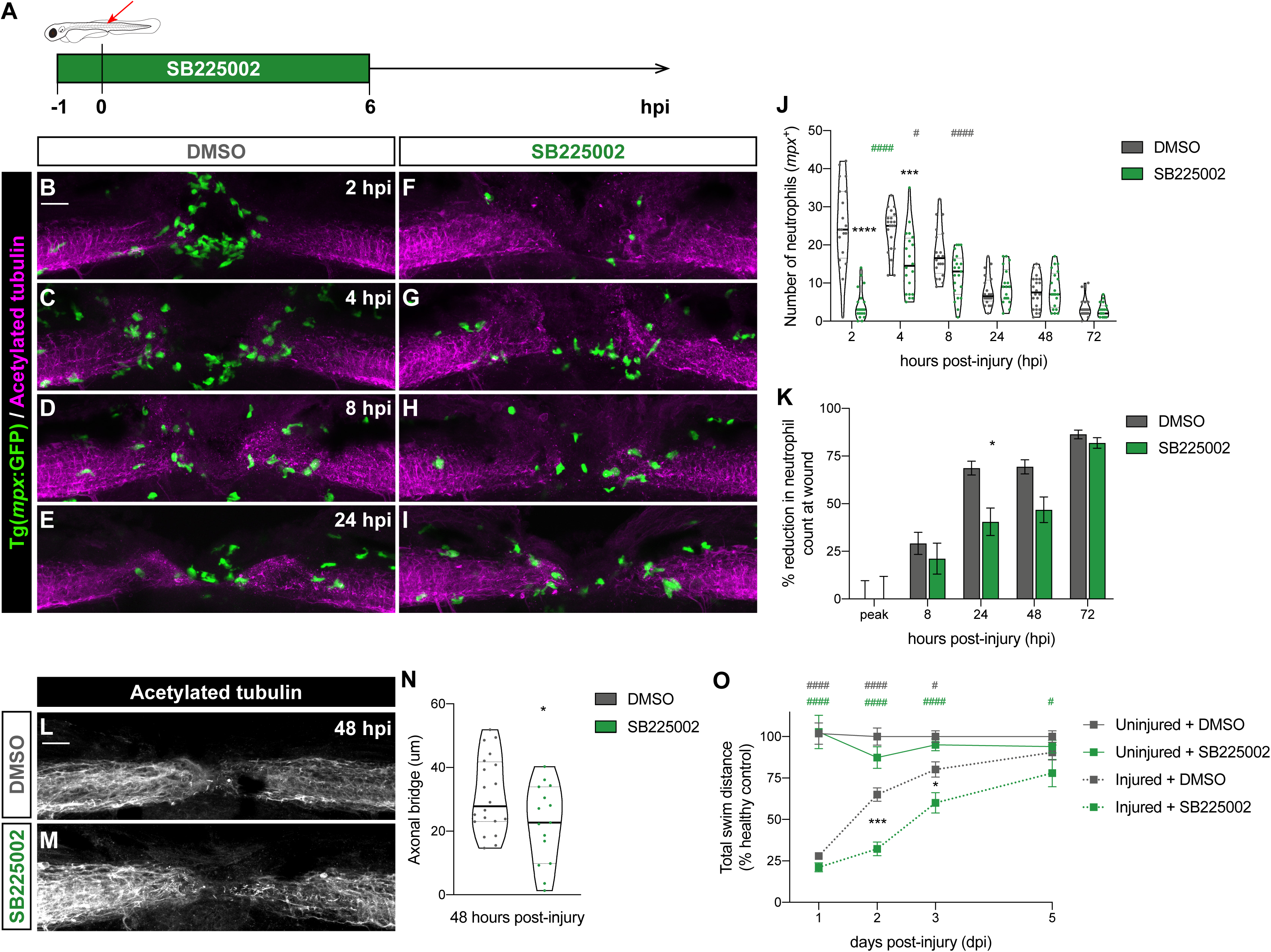
SB225002 treatment delays and attenuates neutrophil recruitment to the injured spinal cord impairing regeneration. **(A)** Scheme depicts the administration of SB225002 in 3 dpf zebrafish larvae, starting one hour before SCI and continuing for the first 6 hours post-injury. **(B-I)** Representative maximal projections reveal neutrophils (green) recruited to the spinal cord (magenta) at 2, 4 and 8 hpi. Scale bar, 30 μm. **(J)** Quantification displays the number of neutrophils at the injured spinal cord, showing a significant delay and reduction in recruitment in SB225002-treated larvae (n = 18 - 22, performed as three independent experiments). Two-way ANOVA followed by Tukey’s multiple comparisons test indicates a significant difference between SB225002- and DMSO-treated larvae (asterisks) at 2 (p < 0.0001) and 4 hpi (p = 0.0004) with no significant difference at 8 (p = 0.0865), 24 ( p > 0.9999), 48 (p > 0.9990) and 72 hpi (p > 0.9999). Statistically significant differences are also observed within each treatment for DMSO-treated larvae (grey pounds) between 4 and 8 hpi (p = 0.0456), and 8 and 24 hpi (p < 0.0001) and for SB225002-treated larvae (green pounds) between 2 and 4 hpi (p < 0.0001). **(K)** Quantification shows the percentage of reduction in neutrophil counts at the wound. Two-way ANOVA followed by Sidak’s multiple comparison test indicates a significant idecrease between SB225002- and DMSO-treated larvae at 24 hpi (p = and no significant difference at the peak (p > 0.9999), 8 (p = 0.9268), 48 (p = 0.0845) and 72 hpi (p = 0.9952). **(L-M)** Representative maximal projections of the spinal cord illustrate the axonal bridge. Scale bar, 30 μm. **(N)** Quantification of the axonal bridge thickness shows a significant decrease in SB225002-treated injured larvae, analyzed via an unpaired two-tailed t-test (p = 0.0349) (n = 15 – 18, performed as three independent experiments). **(O)** Recovery of the swimming capacity expressed as the total swim distance in relationship to the healthy control (n = 29 – 61, performed as three independent experiments). Two-way ANOVA followed by Tukey’s multiple comparison displays significant difference between injured SB225002- and DMSO-treated larvae (asterisks) at 2 (p = 0.0004) and 3 dpi (p = 0.0449); between injured and uninjured DMSO-treated larvae (grey pound) at 1 (p < 0.0001), 2 (p < 0.0001) and 3 dpi (p = 0.0138); and between injured SB225002- and uninjured DMSO-treated larvae (green pound) at 1 (p < 0.0001), 2 (p < 0.0001), 3 (p < 0.0001) and 5 dpi ( p = 0.0278).

Having this impairment in neutrophil dynamics, we wondered whether it would affect the spinal cord regenerative process. Consequently, we assessed this process at both cellular and functional levels by measuring axonal bridging and evaluating swimming capacity recovery in SB225002-treated larvae compared to DMSO controls. At 48 hpi, the SB225002 group showed a 30% decrease in axonal bridge thickness (Figure 2L– N). In addition, when we analyzed the recovery of swimming capacity using a light/dark paradigm, we observed a significant decrease in the total swimming distance of SB225002-treated larvae at 2- and 3-days post-injury (dpi) compared with DMSO-treated controls (Figure 2O). Importantly, this impairment in the recovery of swimming behavior of SB225002 injured larvae was still present at 5 dpi in relation to the DMSO uninjured controls. These results suggest that interfering with neutrophil dynamics through Cxrc1/2 signaling impairs the regenerative process of the spinal cord.

### Accelerating neutrophil clearance from the injured spinal cord improves regeneration

Next, we determined whether altering neutrophil dynamics during the resolution phase affected spinal cord regeneration. The Cxcl12/Cxcr4 signaling axis is involved in retaining neutrophils at sites of tissue damage [35]. AMD3100, a Cxcr4 antagonist, has been used to accelerate neutrophil resolution by promoting reverse migration [35]. Therefore, we administered AMD3100 at 4 hpi, coinciding with the neutrophil recruitment peak, until 24 hpi (Figure 3A). As expected, a 34% reduction in the number of neutrophils at the injury site was observed at 12 hpi in the AMD3100-treated larvae when compared to the DMSO controls (Figure 3B-F), and early neutrophil inflammation resolution was achieved, whereas in DMSO-treated controls, it only occurred at 24 hpi (Figure 3F,G).

**Figure 3:**
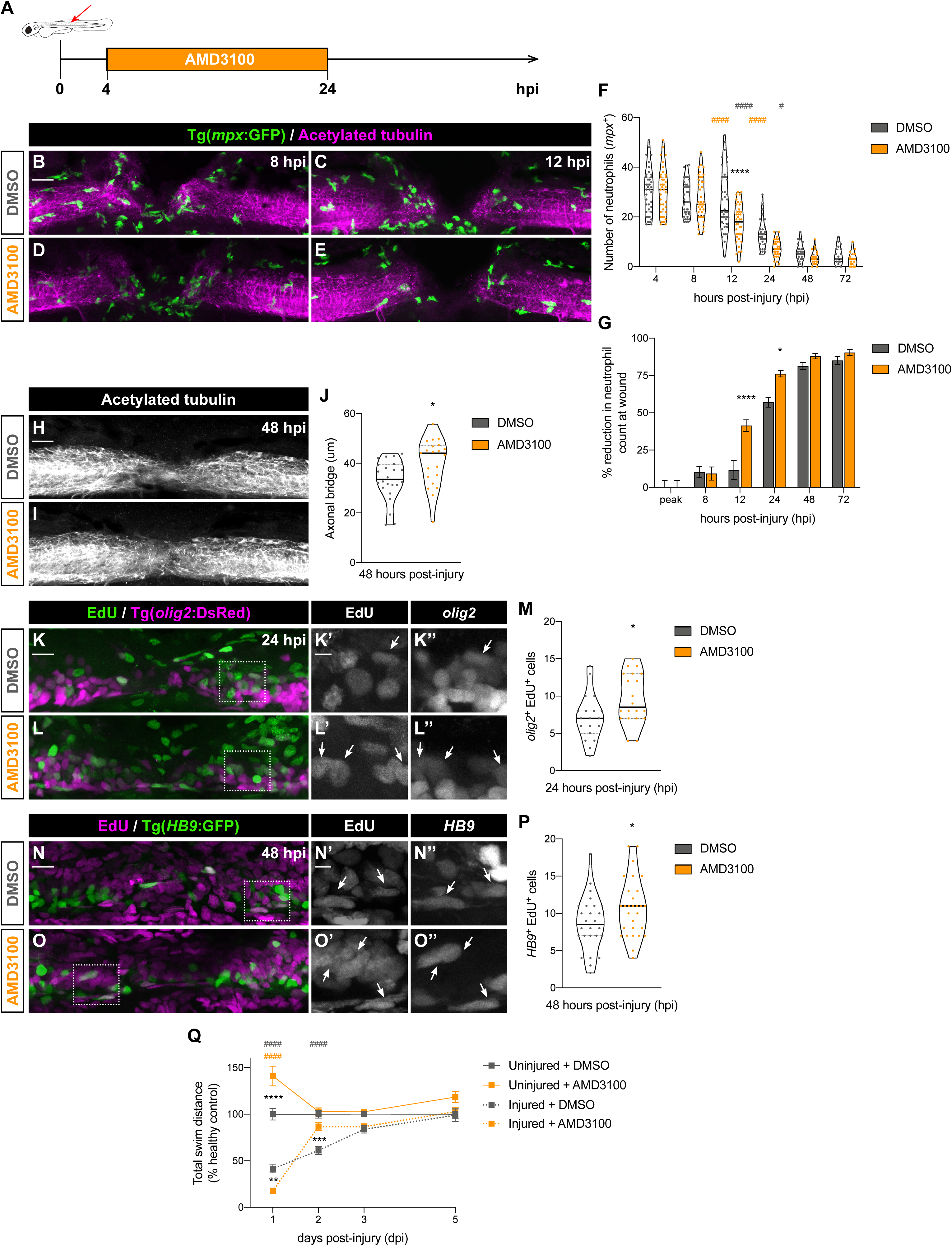
AMD3100 fastens neutrophil clearance from the injured spinal cord spinal cord regeneration. **(A)** Scheme represents the administration of AMD3100 3 dpf larvae, from 4 until 24 hpi. **(B - E)** Representative maximal projections display neutrophils (green) recruited to the injured spinal cord (magenta) at 8 and 12 hpi. Scale bar, 30 μm. **(F)** Quantification of neutrophil numbers at the injury site at 4, 8, 12, 24, 48 and 72 hpi (n = 40 (4, 8 and 12 hpi), 31 (24 hpi), 24 (48 hpi) and 19 (72 hpi), from three independent experiments). Two-way ANOVA followed by Tukey’s multiple comparisons test indicates a significant decrease between AMD3100- and DMSO-treated larvae at 12 hpi (p < 0.0001) (black asterisk) with no significant difference at any other timepoints. Significant differences were also observed within each treatment for DMSO-treated larvae (grey pounds) between 12 and 24 hpi (p < 0.0001) and 24 and 48 hpi (p = 0.0121), and AMD3100-treated larvae (orange pounds) between 8 and 12 hpi (p < 0.0001) and 12 and 24 hpi (p < 0.0001). **(G)** Quantification shows the percentage of reduction in neutrophil counts at the wound. Two-way ANOVA followed by Sidak’s multiple comparison test indicates a significant increase between AMD3100- and DMSO-treated larvae at 12 (p < 0.0001) and 24 hpi (p = 0.0134) and no significant difference at the peak and 8 hpi (p > 0.9999), 48 (p = 0.9282) and 72 hpi (p = 0.9870). **(H, I)** Representative maximal projections of the spinal cord, highlighting the axonal bridge. Scale bar, 30 μm. **(J)** Quantification of the axonal bridge thickness reveals a significant increase between AMD3100- and DMSO-treated injured larvae, analyzed via an unpaired two-tailed t-test (p = 0.0105) (n = 21 – 24, from two independent experiments). **(K, L)** Representative maximal projections of the spinal cord illustrate EdU incorporation in neural progenitors (*olig2^+^* cells). Scale bar, 30 μm. Higher magnification panels (**K’, L’**) and (**K’’, L’’**) indicate regions outlined in rectangular boxes for the Edu and *olig2* channels, respectively. Scale bar, 10 μm. **(M)** Quantification of Edu^+^ *olig2^+^* cells in DMSO- and AMD3100-treated larvae (n = 19 – 20, from three independent experiments) shows a significant difference (p = 0.0213) via an unpaired two-tailed t-test. **(N, O)** Representative maximal projections of the spinal cord exhibit EdU incorporation in motoneurons (*HB9^+^* cells). Scale bar, 30 μm. Higher magnification panels (**N’, O’**) and (**N’’, O’’**) indicate regions outlined in rectangular boxes for the Edu and *HB9* channels, respectively. Scale bar, 10 μm. **(P)** Quantification of Edu^+^ *HB9^+^* cells in DMSO- and AMD3100-treated larvae (n = 19 - 20, from three independent experiments) shows a significant difference (p = 0.0428) via an unpaired two-tailed t-test. **(Q)** Recovery of the swimming capacity, expressed as the total swim distance in relationship to the uninjured control (n = 40 – 71, from three independent experiments). Two-way ANOVA followed by Tukey’s multiple comparison test reveals significant difference between injured AMD3100- and DMSO-treated larvae (black asterisks) at 1 (p = 0.0099) and 2 dpi (p = 0.0002), between uninjured AMD3100- and DMSO-treated larvae (black asterisk) at 1 dpi (p < 0.0001), between injured and uninjured DMSO-treated larvae (grey pounds) at 1 (p < 0.0001) and 2 dpi (p < 0.0001); and between injured AMD3100- and uninjured DMSO-treated larvae (orange pounds) at 1 dpi (p < 0.0001).

Furthermore, we evaluated whether the faster clearance of neutrophils from the injury site in AMD3100-treated larvae would have an impact on the regenerative process. Strikingly, we observed a 21% increase in the thickness of the axonal bridge in the AMD3100 group compared to the DMSO group (Figure 3H-J). Following spinal cord injury, *olig2*-expressing ependymal radial glial cells (ERGs) from the ventral spinal cord give rise to motor neurons [7]. Therefore, we assessed the proliferation of *olig2^+^* cells in our AMD3100-treated larvae in order to unveil the potential contribution of these cells to the thicker axonal bridge. Importantly, a 35% increase in the proliferation of *olig2^+^* progenitor ERGs was observed at 24 hpi in the AMD3100 group (Figure 3K–M). Consequently, a 16% increase in the number of newly generated motor neurons, specifically labelled with *Tg(HB9:GFP)*, was detected in the AMD3100-treated group at 48 hpi (Figure 3N–P). This suggests that the increased proliferation of motor neuron progenitors contributes to the observed increase in axonal bridging.

To determine whether this improvement at the cellular level affected the recovery of swimming capacity, we measured the total swimming distance of AMD3100- and DMSO-treated larvae. Importantly, AMD3100 injured larvae presented a remarkably faster recovery of swimming capacity, showing a similar motility with the uninjured control at 2 dpi, while in DMSO injured larvae, this was only observed at 3 dpi (Figure 3Q). Interestingly, a boost in motility was observed at 1 dpi in uninjured AMD3100-treated larvae, concurrent with the timepoint at which the zebrafish were removed from the drug. AMD3100 or Plerixafor mobilizes hematopoietic stem and progenitor cells (HSPCs) and mature immune cells from the bone marrow to the blood [56] that can result in a doping-like effect. Taken together, these results indicate that increased rate of neutrophil clearance from the injury site promotes proliferation and differentiation of motor neuron progenitors, thereby enhancing spinal cord regeneration.

### Neutrophil dynamics specifically at the injury site determine the spinal cord regenerative outcome

To understand whether the striking spinal cord regenerative phenotypes observed were due to the role of neutrophils at the injury site, we took advantage of a transgenic zebrafish line in which neutrophils express a mutant version of Rac2, whose activity is required for neutrophil motility [57]. In *Tg(mpx:mCherry-2A-Rac2D57N)* zebrafish larvae, neutrophils fail to migrate to the wound and infection sites because of a lack of functional pseudopod formation [20]. As expected, when zebrafish larvae were subjected to spinal cord injury, a clear difference in neutrophil recruitment was observed between *Tg(mpx:mCherry-2A-Rac2D57N)* and *Tg(mpx:mCherry-2A-Rac2)* control larvae (Figure 4A–F). While control Rac2 larvae showed a typical neutrophil infiltration curve, Rac2D57N larvae presented an absence of neutrophils at the injury site at all timepoints (Figure 4G). Next, we evaluated the capacity of Rac2D57N to regenerate the spinal cord by measuring the axonal bridge and analyzing the recovery of the swimming capacity (Figure 4H-K). No differences in any of these parameters were found between Rac2D57N and Rac2, indicating that spinal cord regeneration can occur in the absence of neutrophils at the injury site.

**Figure 4:**
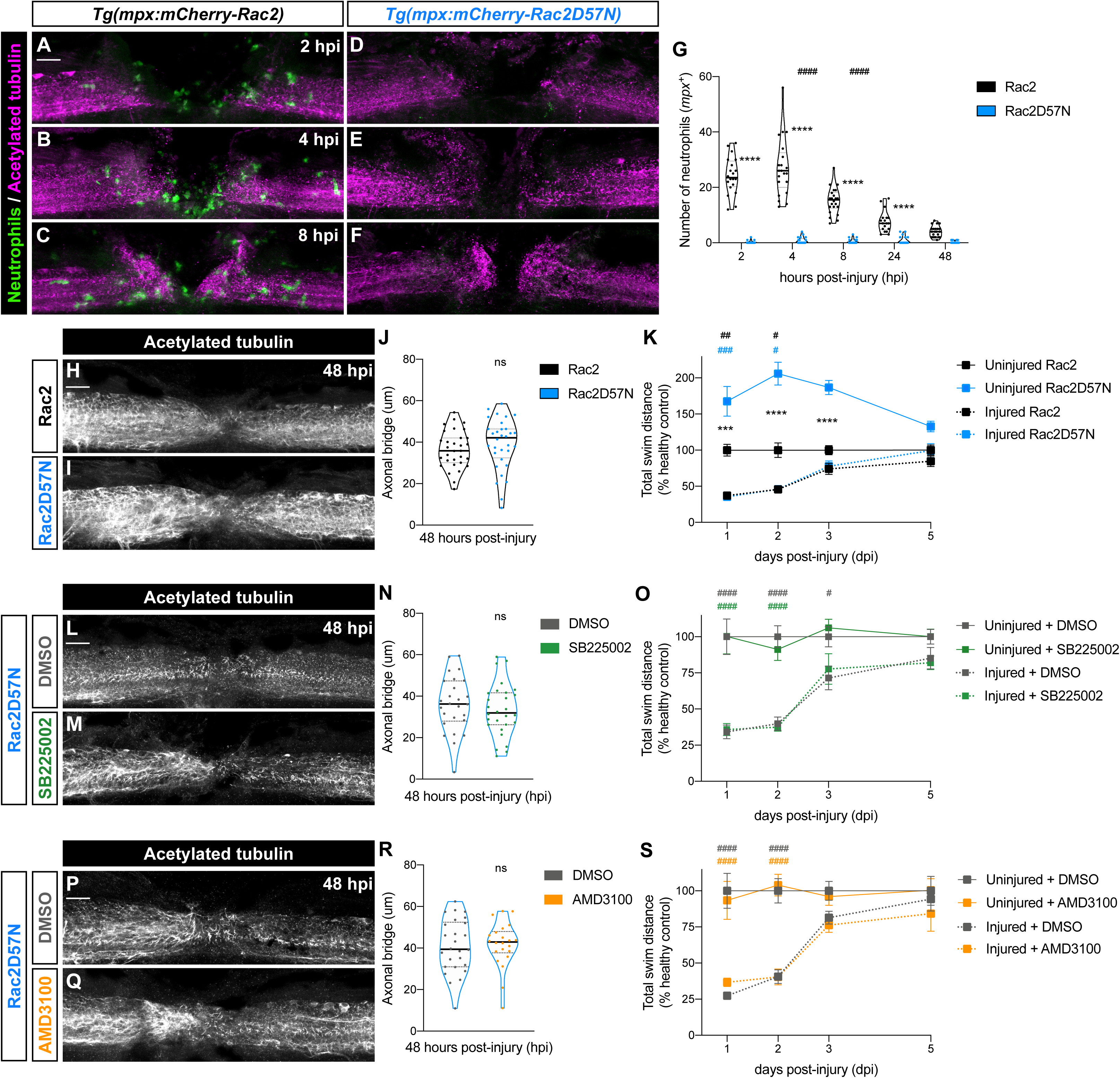
Normal regeneration in Rac2D25N zebrafish larvae following SB225002 or AMD3100 treatment. **(A - F)** Representative maximal projections depict the injury site at 2, 4 and 8 hpi. *Tg(mpx:mCherry-Rac2)* larvae (A – C) show a normal recruitment of neutrophils (green) to the spinal cord (magenta), while no neutrophils were observed in the injured spinal cord of *Tg(mpx:mCherry-Rac2D57N)* larvae (D – F). Scale bar, 30 μm. **(G)** Quantification of *mpx^+^* cells at the injury site demonstrates that neutrophils peak at 2 – 4 hpi in *Tg(mpx:mCherry-Rac2)* larvae. Individual values per larvae are shown (n = 18 - 22 performed in two independent experiments). Two-way ANOVA followed by Tukey’s multiple comparisons test reveals statistically significant differences between 4 and 8 hpi (p < 0.0001) and 8 and 24 hpi (p < 0.0001) in wild-type Rac2 transgenic line. Furthermore, significant differences between wild-type Rac2 and mutant Rac2D57N transgenic lines at 2 (p < 0.0001), 4 (p < 0.0001), 8 (p < 0.0001) and 24 hpi (p < 0.0001) were detected. **(H, I)** Illustrative maximal projections of the spinal cord, highlighting the axonal bridge. Scale bar, 30 μm. **(J)** Quantitative analysis of the axonal bridge thickness indicates no statistically significant difference between wild-type Rac2 and mutant Rac2D57N transgenic larvae, analyzed via unpaired two- tailed t-test (p = 0.3393) (n = 31 – 32, conducted in three independent experiments). **(K)** Recovery of the swimming capacity expressed as the total swim distance (n = 29 – 60, performed in two independent experiments). Two-way ANOVA followed by Tukey’s multiple comparison test reveals no statistically significant difference between injured Rac2D57N and Rac2 larvae at 1, 2, 3 and 5 dpi (p > 0.9999). Statistically significant differences are found between uninjured Rac2D57N and Rac2 larvae (asterisks) at 1 (p = 0.0001), 2 (p < 0.0001) and 3 dpi (p < 0.0001), between injured Rac2D57N and uninjured Rac2 larvae (blue pounds) at 1(p = 0.0009) and 2 dpi (p = 0.0499), and between injured and uninjured Rac2 larvae (black pounds) at 1 dpi (p = 0.0013) and 2 dpi (p = 0.0354). **(L, M)** Representative maximal projections of Rac2D57N injured spinal cord, treated with DMSO or SB225002, illustrate the axonal bridge. Scale bar, 30 μm. **(N)** Quantification of the axonal bridge thickness reveals no statistically significant difference between SB225002- and DMSO-treated Rac2D57N larvae, analyzed via unpaired two tailed t-test (p = 0.4344) (n = 26 – 28, performed in three independent experiments). **(O)** Recovery of the swimming capacity expressed as the total swim distance (n = 27 – 40, performed in two independent experiments). Two-way ANOVA followed by Tukey’s multiple comparison test demonstrates no statistically significant difference between injured SB225002- and DMSO-treated Rac2D57N larvae at 1 (p = 0.9991), 2 (p = 0.9958), 3 (p = 0.9368) and 5 dpi (p = 0.9930). Statistically significant differences are observed between injured and uninjured DMSO-treated larvae (grey pounds) at 1 (p < 0.0001), 2 (p < 0.0001) and 3 dpi (p = 0.0362), as well as between injured SB225002- and uninjured DMSO-treated larvae (green pounds) at 1 (p < 0.0001) and 2 dpi (p < 0.0001). **(P, Q)** Representative maximal projections of Rac2D57N injured spinal cord, treated with DMSO or AMD3100, display the axonal bridge. Scale bar, 30 μm. **(R)** Quantification of the axonal bridge shows no statistically significant difference between AMD3100- and DMSO-treated Rac2D57N larvae, analyzed via unpaired two-tailed t-test (p = 0.8312) (n = 22 – 26, performed in two independent experiments). **(S)** Recovery of the swimming capacity expressed as the total swim distance (n = 24 – 39, performed in three independent experiments). Two-way ANOVA followed by Tukey’s multiple comparison test reveals no statistically significant difference between injured AMD3100- and DMSO-treated larvae at 1 (p = 0.7483), 2 (p > 0.9999), 3 (p = 0.9473) and 5 dpi (p = 0.7990). Statistically significant differences are found between injured and uninjured DMSO-treated larvae (grey pounds) at 1 (p < 0.0001) and 2 dpi (p < 0.0001), as well as between injured AMD3100- and uninjured DMSO-treated larvae (orange pounds) at 1 dpi (p < 0.0001) and 2 dpi (p < 0.0001).

Knowing that Rac2D57N zebrafish larvae were capable of regenerating the spinal cord successfully, we treated them pharmacologically with either SB225002 or AMD3100 and evaluated the regenerative outcome by quantifying the thickness of the axonal bridge and the recovery of swimming capacity. Interestingly, no differences in axonal regrowth were observed following SB225002 or AMD3100 treatment of Rac2D57N larvae (Figure 4L-N, P-R). Accordingly, when the recovery of swimming capacity was measured, no significant differences were observed between SB225002-treated and DMSO-treated (Figure 4O) or between AMD3100-treated and DMSO-treated Rac2D57N larvae (Figure 4S). These observations indicate that the impairment or improvement in spinal cord regeneration observed following SB225002 or AMD3100 treatment, respectively, is only detected when neutrophils are present at the site of injury. This suggests that altering neutrophil dynamics through the Cxcr1/2 and Cxcr4 signaling pathways has an essential effect on the regenerative outcome.

### Cxcr4 inhibition enhances neutrophil activation at the injury site

Owing to its translational relevance, we sought to determine the biological mechanisms promoted by neutrophils that improve spinal cord regeneration upon Cxcr4 inhibition. Therefore, we sorted *mpx:GFP^+^* cells from the injury site of lesioned larvae and from the same region of age-matched uninjured controls, and performed RNA sequencing (RNA-seq) analysis at 8 hpi (Figure 5A). Importantly, the obtained dataset validated the high purity and specificity of the sorted samples showing high expression of neutrophil-specific genes (*mpx*, *lyz*) and immune markers (*mmp9*, *mmp13a*, *coro1a*, *ptpn6*), and absence of macrophage- and microglia-specific genes (*mpeg1.1*, *mfap4*, *irf8*, *marco*, *csf1ra*, *p2ry12*) (Figure S1A). In addition, PCA comparing all RNA-seq samples showed that the biological replicates for each condition clustered together, highlighting their similar transcription profiles (Figure S1B). Moreover, the greatest variation in gene expression was due to injury (PC1), while the second most variation was due to treatment (PC2).

**Figure 5:**
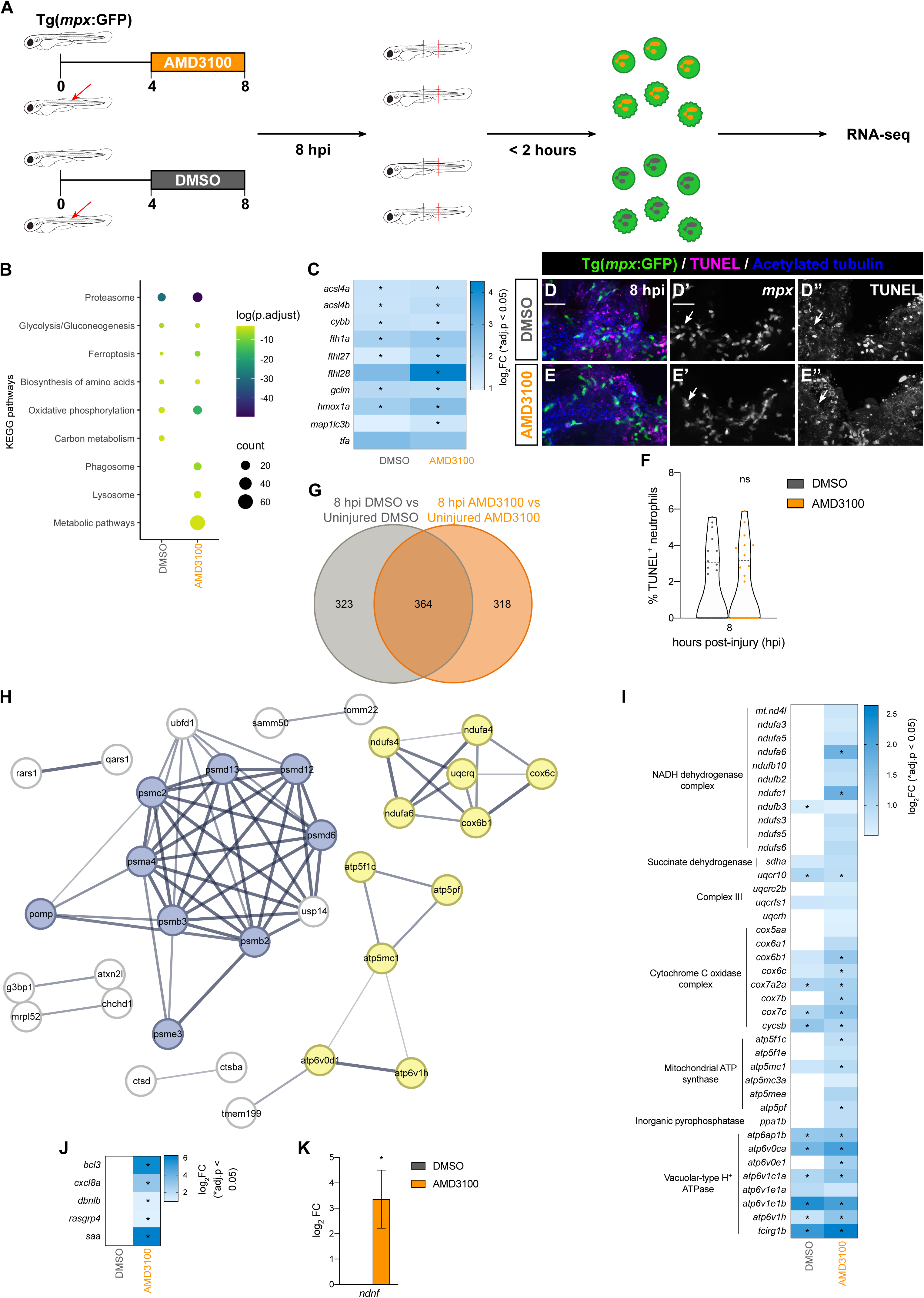
Injury responsive neutrophils exhibit an enhanced activation state upon AMD3100 treatment. **(A)** Scheme illustrating the experimental design. **(B)** KEGG pathways analysis of DEGs (p.adjust < 0.05) in the 8 hpi DMSO vs uninjured DMSO (DMSO) and in the 8 hpi AMD3100 vs uninjured AMD3100 (AMD3100) comparisons. Circle color and size correspond to the log (p. adjust) value and the number of genes, respectively. **(C)** Heatmap analysis of DEGs related to ferroptosis in the DMSO and AMD3100 comparisons. Log_2_ fold change is displayed as blue color gradient when genes present a p value < 0.05. Stars indicate statistical significance (p.adjust < 0.05). **(D, E)** Representative maximal projection illustrating TUNEL+ neutrophils in the injured spinal cord at 8 hpi in DMSO and AMD3100-treated larvae. Scale bar, 30 μm. Individual panels for *mpx* (**D’, E’**) and TUNEL staining (**D’’, E’’**) indicate regions outlined in the rectangular boxes. **(F)** Quantification of the percentage of neutrophils positive for TUNEL at 8 hpi in DMSO and AMD3100-treated larvae (n = 28-29, performed as two independent experiments). Unpaired two-tailed t-test shows no statistically significant difference between DMSO and AMD3100-treated larvae (p = 0.8949). **(G)** Venn diagram depicting DEGs (p adjust < 0.05) in 8 hpi DMSO compared to uninjured DMSO and in 8 hpi AMD3100 compared to uninjured AMD3100. **(H)** Network analysis of AMD3100-specific DEGs related to proteasome (blue) and oxidative phosphorylation (yellow). Protein-protein interaction enrichment p = 8.83e-13. **(I)** Heatmap analysis of DEGs related to oxidative phosphorylation in the 8 hpi DMSO vs uninjured DMSO (DMSO) and 8 hpi AMD3100 vs uninjured AMD3100 (AMD3100) comparisons. Log_2_ fold change is displayed as blue color gradient when genes present a p value < 0.05. Stars indicate statistical significance (p.adjust < 0.05). **(J)** Heatmap analysis depicting DEGs related to activated neutrophils in the 8 hpi DMSO vs uninjured DMSO (DMSO) and 8 hpi AMD3100 vs uninjured AMD3100 (AMD3100) comparisons. Log_2_ fold change is displayed as blue color gradient, and stars indicate statistical significance (p.adjust < 0.05). **(K)** Bar graph illustrating the upregulated expression of neuron-derived neurotrophic factor in the AMD3100 (8 hpi vs. uninjured) comparison.

Next, we investigated the neutrophil response to spinal cord injury by retrieving the statistically significant differentially expressed genes (DEGs) from the 8 hpi vs. uninjured comparisons following AMD3100 or DMSO treatment. By exploring the enrichment in Kyoto Encyclopedia of Genes and Genomes (KEGG) pathways, we identified that glycolysis/gluconeogenesis and biosynthesis of amino acids, which are essential pathways for neutrophils to exert their function and promote their survival at the injury site, were similarly enhanced in both conditions [58] (Figure 5B). In addition, ferroptosis, a regulated cell death characterized by iron-dependent accumulation of lipid peroxides, was also upregulated in both conditions (Figure 5B), together with a significant increase of genes involved in iron metabolism (*fth1a*, *fthl27*, *fthl28*, *hmox1a*, *tfa*) and lipid peroxidation (*acsl4a*, *acsl4b*, *cybb*) (Figure 5C). These findings correlate with the observation of 1-2% of neutrophils that show signs of irreversible cell death at the injury site (Figure 5D-F). This suggests that following spinal cord injury, a small fraction of neutrophils die at the injury site by ferroptosis.

To determine the underlying causes of the enhanced regenerative capacity induced by neutrophils following Cxcr4 inhibition, we compared the statistically significant DEGs triggered by the injury and identified the AMD3100-specific gene set (Figure 5G). Protein network analysis showed a statistically significant enrichment in protein-protein interactions within this gene set, with proteasome and oxidative phosphorylation (OXPHOS) being the top two enriched pathways (Figure 5H). Within the OXPHOS pathway, several genes involved in the mitochondrial electron transport chain and vacuolar-type H^+^ ATPase were upregulated (Figure 5I). Neutrophils use glycolysis as the main metabolic pathway to generate the necessary ATP to meet their energy needs [58, 59], whereas mitochondrial OXPHOS is an important source of mitochondria reactive oxygen species (mROS). Moreover, KEGG pathway enrichment analysis revealed that phagosome, lysosome, and metabolic pathways were specifically enriched in the AMD3100 group (Figure 5B). Consistent with the idea that the upregulation of these pathways, together with vacuolar ATPases, indicates enhanced neutrophil activation [60], we also identified genes involved in the regulation of immune response (*cxcl8a*, *saa*) and neutrophil activation (*bcl3*, *dbnlb*, *rasgrp4*) that were specifically upregulated in the AMD3100 group (Figure 5J). Finally, a strong upregulation of the neuron-derived neurotrophic factor (*ndnf)* was observed in our AMD3100 injury-responsive neutrophils, which was completely absent in the DMSO group (Figure 5K). Taken together, our RNA-seq data suggest that AMD3100 treatment enhances neutrophil activation and promotes the expression of growth factors prior to initiating the resolution of neutrophil inflammation process.

### Neutrophil enhanced activation via Cxcr4 inhibition induces a transitory pro-inflammatory response in macrophage/microglia

Given the transcriptomic signatures found in neutrophils treated with AMD3100, which suggested an enhanced neutrophil activity at the injury site, we hypothesized that this would affect other immune populations. Therefore, we isolated RNA from dissected larval trunks containing the injury site and performed qPCR analysis. At 8 hpi, we observed an upregulation of *tnf-α* and *il-1*β, pro-inflammatory cytokines, in the tissues of AMD3100-treated larvae compared to the DMSO group (Figure 6A). These pro-inflammatory cytokines play essential roles at the beginning of the spinal cord regenerative process, but their sustained expression can be detrimental [9].

**Figure 6:**
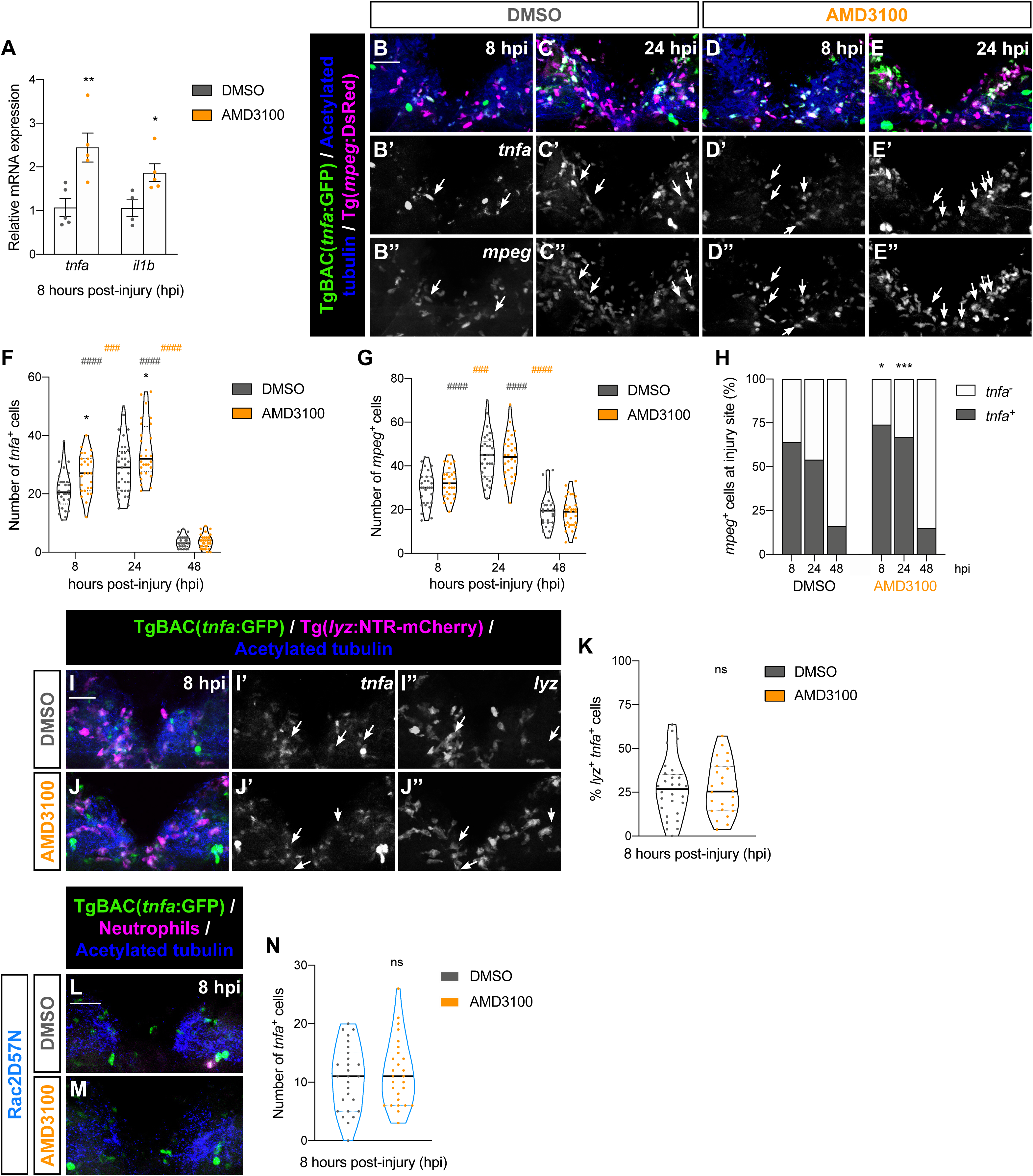
Enhanced proportion of *mpeg^+^ tnfa^+^* cells following AMD3100 treatment. **(A)** Relative gene expression, measured by qPCR, of proinflammatory cytokines. Each dot represents a biological replicate obtained from a pool of thirty trunks and each bar depicts the mean ± SEM. Unpaired two-tailed t-test shows a significant upregulation between DMSO and AMD3100-treated larvae at 8 hpi for *tnfa* (p = 0.0076) and *il1b* (p = 0.0257). **(B - E)** Representative maximal projections display *mpeg^+^* cells and *tnfa^+^* cells in the injured spinal cord at 8 and 24 hpi in DMSO and AMD3100-treated larvae. Scale bar, 30 μm. Individual panels for *tnfa* (**B’ – E’**) and *mpeg* (**B’’ - E’’**) where double-positive cells are indicated by arrows. **(F)** Quantification of the number of *tnfa^+^* cells in the injured spinal cord at 8, 24 and 48 hpi in DMSO and AMD3100-treated larvae (n = 26 - 33, conducted in three independent experiments). Two-way ANOVA followed by Tukey’s multiple comparison test reveals a statistically significant difference between AMD3100- and DMSO-treated larvae (asterisk) at 8 hpi (p = 0.0181) and 24 hpi (p = 0.0132), with no statistically significant difference at 48 hpi (p > 0.9999). Additionally, within the DMSO-treated larvae (grey pounds), significant differences were observed between 8 and 24 hpi (p < 0.0001) and 24 and 48 hpi (p < 0.0001). Within the AMD3100-treated group (orange pounds), differences were observed between 8 and 24 hpi (p = 0.0003) and 24 and 48 hpi (p > 0.0001). **(G)** Quantification of *mpeg^+^* cell number in the injured spinal cord at 8, 24 and 48 hpi in DMSO- and AMD3100-treated larvae (n = 26 - 33, performed in three independent experiments). Two-way ANOVA followed by Tukey’s multiple comparison test exhibits statistically significant difference between 8 and 24 hpi and 24 and 48 hpi (p < 0.0001 for both DMSO and AMD3100-treated larvae in both timepoint comparisons). **(H)** Quantification of the percentage of *mpeg^+^ tnfa^+^* cells in the injured spinal cord at 8, 24 and 48 hpi in DMSO and AMD3100-treated larvae (n = 26 – 33, conducted in three independent experiments). Two-way ANOVA followed by Sidak’s multiple comparisons test shows a statistically significant increase between AMD3100 and DMSO-treated larvae at 8 hpi (p = 0.0283) and 24 hpi (p = 0.0005), with no statistically significant difference at 48 hpi (p = 0.9250). **(I, J)** Representative maximal projection illustrate neutrophils (*lyz^+^* cells) and *tnfa^+^* cells in the injured spinal cord at 8 hpi in DMSO and AMD3100-treated larvae. Scale bar, 30 μm. Arrows in individual panels for *tnfa* **(I’, J’)** and *lyz* **(I’’, J’’)** highlight double-positive cells. **(K)** Quantification of the percentage of *lyz^+^ tnfa^+^* cells in the injured spinal cord at 8 hpi in DMSO and AMD3100-treated larvae (n = 24 – 28, performed in two independent experiments). Unpaired two-tailed t-test shows no statistically significant difference between DMSO and AMD3100-treated larvae at 8 hpi (p = 0.9504). **(L, M)** Representative maximal projections illustrate *tnfa^+^* cells in the injured spinal cord of Rac2D57N larvae, treated with DMSO or AMD3100, at 8 hpi. Scale bar, 30 μm. **(N)** Quantification of the number of *tnfa^+^* cells in the injured spinal cord at 8 hpi in DMSO and AMD3100-Rac2D57N larvae (n = 27, conducted in two independent experiments). Unpaired two-tailed t-test shows no statistically significant difference between DMSO and AMD3100-treated larvae at 8 hpi (p = 0.5767).

To understand the origin of the increased *tnf-α* expression, we employed the *Tg(mpeg1.1:mCherryF) TgBAC(tnfa:GFP)* double transgenic line to label macrophage/microglia, and *tnf-α*-expressing cells, respectively [25, 26] (Figure 6B–E). In agreement with the qPCR results, 24% and 20% increases in *tnf-α*-expressing cells were detected after AMD3100 treatment at 8 and 24 hpi, respectively, in the spinal cord (Figure 6F). This enhanced presence of *tnf-α*-expressing cells did not alter *tnf-α* dynamics during the regenerative process since an expected increase was observed between 8 and 24 hpi, which sharply decreased by 48 hpi [9]. Moreover, although no difference was observed in the absolute number of *mpeg^+^* cells in the injured spinal cord between the AMD3100 and DMSO groups (Figure 6G), 16% and 23% increases in the proportion of *mpeg^+^ tnf-α^+^* cells were detected at 8 and 24 hpi, respectively (Figure 6H). In addition, when neutrophils were evaluated, no increase in the proportion of *tnf-α*-expressing neutrophils was detected (Figure 6I-K). This suggests that enhancing neutrophil activation transcriptomic signatures at the injury site via Cxcr4 inhibition contributes to early increased proportion of pro-inflammatory macrophage/microglia at the injury site. Interestingly, *tnf-α*-expressing macrophages were shown to promote ERG proliferation and regenerative neurogenesis in the zebrafish spinal cord [13]. Therefore, this transient early increase in the proportion of *tnf-α*-expressing macrophage/microglia could explain the enhanced spinal cord regeneration observed in AMD3100-treated larvae.

Furthermore, to confirm that this increase in the proportion of pro-inflammatory macrophage/microglia was derived from modulation of neutrophil activity, we treated Rac2D57N zebrafish larvae, which lack neutrophils at site of injury, with AMD3100 and quantified the number of *tnf-α*-expressing cells (Figure 6L–N). As expected, when neutrophils were not present at the injury site, no difference in the number of *tnf-α^+^* cells was detected (Figure 6N). This result confirmed our initial hypothesis, demonstrating that modulation of neutrophil activity affects other immune populations at the injury site, thus contributing to the accelerated spinal cord regeneration.

### Cxcr1/2-inhibited neutrophils at the injury site impair spinal cord regeneration

Knowing the effects of neutrophils on other immune populations, we attempted to identify the underlying cause of the spinal cord regenerative impairment observed when neutrophil recruitment to the injury site was altered (Figure 2). To evaluate macrophage/microglial dynamics, we used the *Tg(mpeg1.1:mCherryF)* zebrafish line. Interestingly, while the DMSO-treated group showed a normal infiltration and resolution curve of *mpeg^+^* cells at the injury site, peaking at 24 hpi, in the SB225002-treated larvae, a strong reduction in *mpeg^+^* cell numbers (53% at 8 hpi, 49% at 24 hpi, and 30% at 48 hpi) and a flat infiltration pattern were observed (Figure 7A-E). As previously shown, SB225002 is a specific Cxcr1/2 antagonist that inhibits neutrophil recruitment to the injury site in a broad variety of tissues [45, 52, 54]. However, neutrophils are not the only immune cells expressing *cxcr1/*2, and macrophages have recently been reported to express a *cxcr1-like* receptor in zebrafish larvae [61]. Therefore, to understand whether SB225002 treatment was responsible for the reduction in *mpeg^+^* cell numbers at the injury site, we treated Rac2 and Rac2D57N larvae with DMSO and SB225002, and quantified the number of *mpeg^+^* cells using *Tg(mpeg:EGFP)*. Notably, a similar significant decrease in *mpeg^+^* cells at the injury site in SB225002-treated larvae compared to the DMSO control group was observed in Rac2 (69% at 8 hpi and 81% at 24 hpi) and Rac2D57N (57% at 8 hpi and 60% at 24 hpi) zebrafish larvae (Figure 7H-P). This demonstrates that the reduction in *mpeg^+^* cells at the injury site occurs independently of the presence of neutrophils, being a direct effect of SB225002 treatment. Strikingly, when *mpeg^+^* dynamics over time were analyzed, the expected peak at 24 hpi was only observed in DMSO-treated Rac2 and Rac2D57N larvae and in SB225002-treated Rac2D57N larvae (Figure 7P). Rac2 larvae treated with SB225002 did not show a significant increase in *mpeg^+^* cells at the injury site, which is in accordance with our previous observations (Figure 7E). This indicates that inhibiting Cxcr1/2 signaling axis in neutrophils at the injury site decreases the presence of *mpeg^+^* cells, thereby exerting some control over the recruitment of these immune populations to the site of injury.

**Figure 7:**
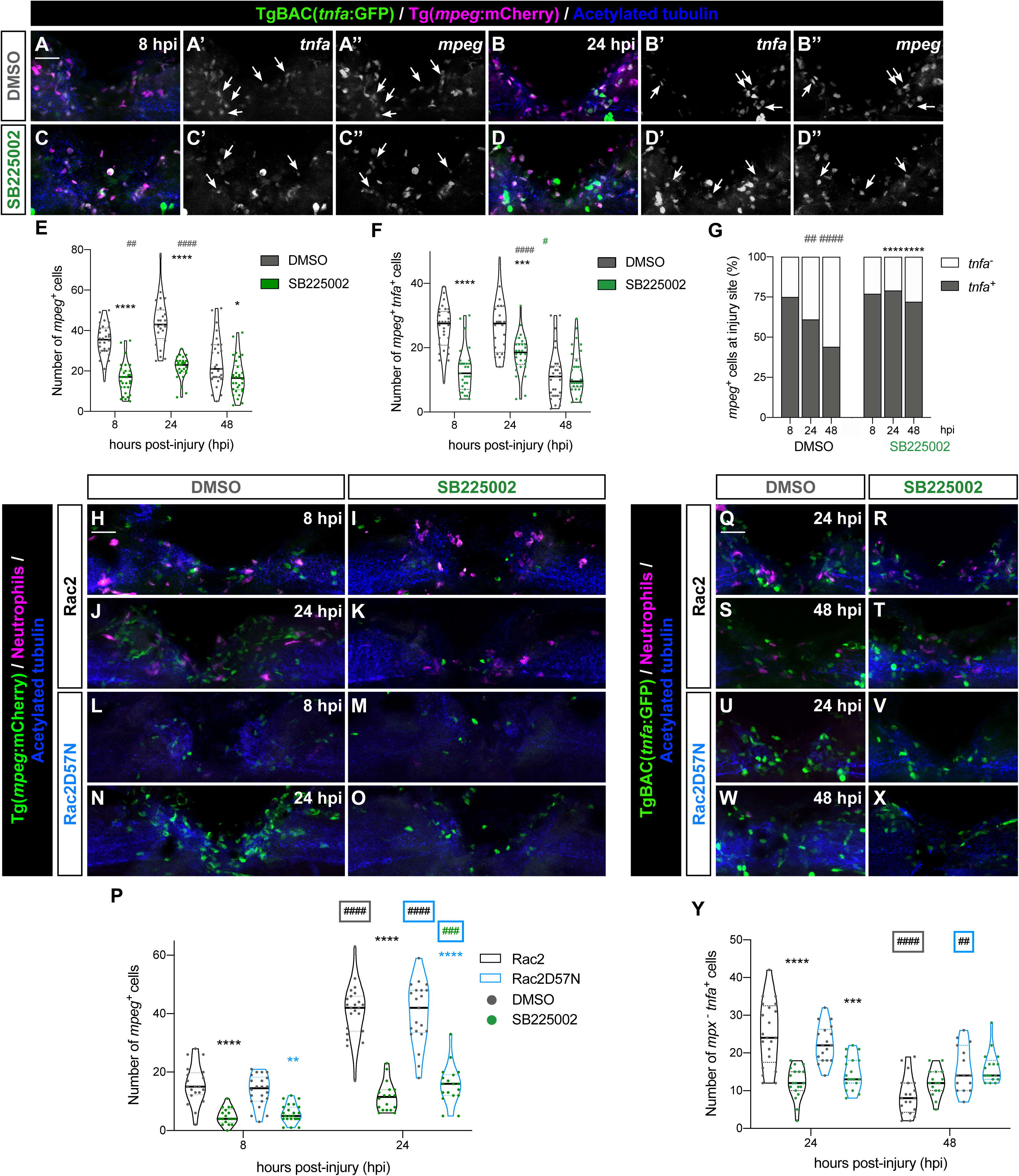
SB225002-treated neutrophils attenuate *mpeg^+^* presence at the injury site. **(A – D)** Representative maximal projections displaying *mpeg^+^* and *tnfa^+^* cells in the injured spinal cord at 8 and 24 hpi in DMSO- and SB225002-treated larvae. Scale bar, 30 μm. Individual panels for *tnfa* (**A’ - D’**) and *mpeg* (**A’’ - D’’**) highlight double-positive cells. **(E)** Quantification of *mpeg^+^* cells in the injured spinal cord (n = 24 - 31, performed as two independent experiments). Two-way ANOVA followed by Tukey’s multiple comparison test revealed a statistically significant difference between DMSO- and SB225002-treated larvae at 8 (p < 0.0001), 24 (p < 0.0001) and 48 hpi (p = 0.0315). Additionally, significant differences were observed in DMSO-treated larvae between 8 and 24 hpi (p = 0.0079) and between 24 and 48 hpi (p < 0.0001) (grey pounds), with no significant difference in SB225002-treated larvae between 8 and 24 hpi (p = 0.2161) and 24 and 48 hpi (p = 0.3950). **(F)** Quantification of the number of *mpeg^+^ tnfa^+^* cells in the injured spinal cord (n = 24 - 31, performed as two independent experiments). Two-way ANOVA followed by Tukey’s multiple comparison test showed a statistically significant decrease between DMSO- and SB225002-treated larvae at 8 (p < 0.0001) and 24 hpi (p = 0.002), with no significant difference at 48 hpi (p = 0.9991). **(G)** Quantification of the proportion of *mpeg^+^ tnfa^+^* cells over the total *mpeg^+^* population (n = 24 - 31, performed as two independent experiments). Two-way ANOVA followed by Tukey’s multiple comparison test showing statistically significant increase between DMSO- and SB225002-treated larvae at 24 (p < 0.0001) and 48 hpi (p < 0.0001). Additionally, a significant decrease was observed in the DMSO-treated group between 8 and 24 hpi (p = 0.0014) and 24 and 48 hpi (p < 0.0001). **(H – O)** Representative maximal projections illustrate *mpeg^+^* cells in transgenic Rac2 and Rac2D57N DMSO- and SB225002-treated larvae at 8 and 24 hpi. Scale bar, 30 μm. **(P)** Quantification of *mpeg^+^* cells in the injured spinal cord in Rac2 and Rac2D57N larvae (n = 18 - 23, performed as two independent experiments). Two-way ANOVA followed by Tukey’s multiple comparison test showed a significant decrease between DMSO- and SB225002-treated larvae at 8 (p < 0.0001) and 24 hpi (p < 0.0001) in Rac2 larvae and at 8 (p = 0.0039) and 24 hpi (p < 0.0001) in Rac2D57N larvae. Additionally, significant increases were observed between 8 and 24 hpi in DMSO- Rac2 (p < 0.0001) and Rac2D57N (p < 0.0001) and SB225002- Rac2D57N treated larvae (p = 0.0002). No statistically significant difference in SB225002 Rac2 larvae between 8 and 24 hpi was observed (p = 0.0707). **(Q – X)** Representative maximal projections exhibit *tnfa^+^* cells in transgenic Rac2 and Rac2D57N. Scale bar, 30 μm. **(Y)** Quantification of *mpx^-^ tnfa^+^* cells in the injured spinal cord in Rac2 and Rac2D57N (n = 15 - 21, performed as three independent experiments). Two-way ANOVA followed by Tukey’s multiple comparison test showed a statistically significant difference at 24 hpi between DMSO and SB225002-treated Rac2 (p < 0.0001) and Rac2D57N (p = 0.0004) larvae, with no significant difference at 48 hpi in Rac2 (p = 0.5070) and Rac2D57N (p > 0.9999) larvae. Additionally, significant decreases from 24 to 48 hpi were only observed in DMSO-treated Rac2 (p < 0.0001) and Rac2D57N (p = 0.0098) larvae.

Next, to determine whether *mpeg^+^* cells presented an abnormal pro-inflammatory profile after SB225002 treatment, we evaluated the number of *tnf-α*-expressing *mpeg^+^* cells using the *TgBAC(tnfa:GFP)* zebrafish line (Figure 7A-D, F). Interestingly, an increased proportion of *mpeg^+^ tnf-α^+^* cells sustained over time, was observed in SB225002-treated larvae compared to the DMSO group (Figure 7G). Inflammation has to be tightly controlled to lead to successful resolution and regeneration [62], and sustained elevated Tnf-α levels have been associated with impairments in retinal regeneration in zebrafish [63]. Therefore, we wondered whether this increased proportion of pro-inflammatory *mpeg^+^* cells was the cause of the impaired regeneration (Figure 2L-O). During spinal cord regeneration, *tnf-α* is mainly expressed by immune cells [9], and the immune system of zebrafish larvae consists of only innate populations [64]. To answer this question, we first treated Rac2 and Rac2D57N zebrafish larvae with SB225002 and DMSO and quantified the number of *mpx^-^ tnf-α^+^* cells (Figure 7Q-Y). Interestingly, similar cellular dynamics were observed in SB225002 Rac2- and Rac2D57N-treated larvae when compared to the DMSO groups. A significant decrease in absolute *mpx^-^ tnf-α^+^* cell numbers at 24 hpi, which normalized by 48 hpi, was observed in Rac2 (40%) and Rac2D57N (35%), following a similar trend to that previously observed for *mpeg^+^ tnf-α^+^* cell numbers in wild-type larvae (Figure 7F). This suggests that the sustained increase in the proportion of *mpeg^+^ tnf-α^+^* cells is a direct effect of SB225002 treatment and is not an effect caused by the inhibition of Cxcr1/2 signaling in the neutrophils at the injury site. In addition, we sought to rescue the regenerative impairment by decreasing the enhanced pro-inflammatory levels after SB225002 treatment using Metformin, which modulates inflammation by downregulation of *tnf-α* [32]. Metformin treatment from 24 hpi to 48 hpi, significantly lowered the proportion of *mpeg^+^ tnf-α^+^* cells at the injury site at 48 hpi by 26% but this reduction did not significantly improve the axonal bridging (Figure S6). Taken together, our data suggest that a reduction in the number of *mpeg^+^* cells at the injury site, in combination with a prolonged increase in their pro-inflammatory profile, does not fully explain the observed impairment in axonal regeneration, at least in our experimental settings. In addition, our results suggest that SB225002-treated neutrophils recruited to the injury site may present an inflammatory profile that not only reduces the presence of *mpeg^+^* cells but is also detrimental to the spinal cord regenerative process. Future investigations into the effects of blocking Cxcr1/2 signaling in zebrafish neutrophils will help elucidate the mechanisms underlying this regenerative impairment.

## DISCUSSION

This study demonstrates that altering neutrophil dynamics at the injury site through Cxcr1/2 and Cxcr4 signaling inhibition has a profound effect on spinal cord regeneration in zebrafish. Delaying neutrophil infiltration via Cxcr1/2 inhibition impairs the regenerative process, whereas promoting early neutrophil inflammation resolution through Cxcr4 inhibition accelerates it. Mining the neutrophil-specific RNA-seq data generated in this study, allowed us to reveal how blocking Cxcr4 signaling alters the transcription profile towards an enhanced activation state. This change in the neutrophil inflammatory profile translates into a transient increased expression of *tnf-α* in macrophage/microglia. Conversely, blocking Cxcr1/2 signaling indicates a detrimental inflammatory profile in neutrophils that negatively affects the accumulation of macrophage/microglia at the injury site.

Our studies complement and extend previous research demonstrating the immune heterogeneity of neutrophils and their role in shaping the immune response to injury. CXCR2 and CXCR4 signaling, in addition to their role in neutrophil chemotaxis, are crucial in balancing the neutrophil response between defensive and protective states [65]. In line with this, compounds promoting the resolution of neutrophil inflammation have been proposed as potential therapeutic targets [21, 66]. Supporting this concept, recent research has shown that promoting neutrophil reverse migration increases *tnf-α^+^* macrophages and enhances cardiac regeneration in zebrafish [67]. Here, we report a boost in spinal cord regeneration derived from promoting early neutrophil clearance through the inhibition of Cxcr4 signaling. Our neutrophil-specific RNA-seq data reveals that blocking this signaling axis induces upregulation of the mitochondrial oxidative phosphorylation (OXPHOS), proteasome, phagosome, lysosome, and metabolic pathways. Mitochondrial OXPHOS is involved in regulating respiratory burst, development, and migration [68, 69], whereas phagosome and proteasome pathways are intrinsic to neutrophil activity. Therefore, our results suggest that inhibiting Cxcr4 signaling increases neutrophil cellular activity, switching them towards a more “defensive” state and potentially a pro-resolution phenotype. Additionally, our data demonstrate that this alteration in the neutrophil immune profile is associated with an early and transient increase in the polarization state of macrophage/microglia towards a *tnf-α^+^* phenotype. This priming of macrophage inflammatory state by neutrophils aligns with previous reports in an influenza model, in atherosclerosis, and especially in an acute liver injury mouse model, where neutrophil-derived ROS, triggers monocyte differentiation into a pro-repair state to promote tissue resolution [70–72]. This example could easily explain our boost in spinal cord regeneration, considering that *tnf-α* expressed by macrophages increases neurogenesis in zebrafish [13]. In addition to *tnf-α* expressed by macrophages, it is expected that additional neurogenic factors might contribute to spinal cord regeneration. We observe an increase in the expression of neural growth factor *ndnf* in neutrophils primed with Cxcr4 inhibition. *Ndnf*, is an evolutionarily conserved growth factor expressed in neural tissues in the brain and spinal cord during embryonic development in zebrafish and mice [73, 74]. The capacity of neutrophils to secrete growth factors to promote central nervous system repair has previously been reported [15], and could also explain the improved regeneration phenotype. Further investigations analyzing whether neutrophils, directly or indirectly (through their action on macrophage/microglia), improve spinal cord regeneration will uncover key new players with regenerative potential. Taken together, our results demonstrate that blocking Cxcr4 signaling in neutrophils affects their inflammatory state and behavior, impacting on different cell types within the spinal cord microenvironment.

Our data illustrates that inhibiting Cxcr1/2 signaling results in a delayed and diminished infiltration of neutrophils to the spinal cord injury site that was associated with a late resolution of neutrophil inflammation. This observation aligns with previous work showing that neutrophil migratory behavior towards the wound site was also altered upon inhibition of this chemoattractant signaling axis [51]. Neutrophils express an extensive variety of receptors that can sense pro-inflammatory mediators and modulate neutrophil polarization and recruitment [48, 75]. As it has been previously suggested [51], it is possible that neutrophil subtypes express distinct receptors and are recruited to the injury site through different chemotactic pathways, which can ultimately regulate their polarization and inflammatory profiles impacting the overall inflammatory response, resolution, and wound healing processes. In addition, neutrophil-secreted molecules act as chemoattractants regulating the extravasation of inflammatory monocytes [76]. This aligns with our data, which shows that blocking Cxcr1/2 signaling in neutrophils ablates macrophage/microglia infiltration peak. Interestingly, a similar effect on *mpeg^+^* cell dynamics has been recently reported in zebrafish with an impact on heart repair [45]. Further studies analyzing the transcriptional changes induced by the blockage of Cxcr1/2 signaling in zebrafish neutrophils will help to understand the detrimental effects caused in tissue regeneration. In addition, it would be interesting to compare this expression profile with mammalian datasets, where the regenerative capacity is very low, to evaluate the presence of common biological pathways. Moreover, our experimental setup reveals a previously undescribed side effect of SB225002 drug on the *mpeg^+^* population. This suggests that zebrafish *mpeg^+^* cells have Cxcr1/2 receptors similar to mammals. Importantly, this side effect does not affect the repair process, underscoring the importance of Cxcr1/2 signaling in neutrophils for tissue regeneration.

Contrary to the classical understanding that neutrophil inflammation resolution occurs through apoptosis, followed by macrophage phagocytosis, recent research has identified neutrophil reverse migration as an evolutionarily conserved mechanism for local inflammation resolution [46, 77, 78]. However, the biological role of this process, its regulation, and the fate of neutrophils that undergo reverse migration remain unknown. Our results indicate that the majority of neutrophils recruited to the injured spinal cord in zebrafish larvae undergo reverse migration to multiple tissues throughout the body. These reverse-migrated neutrophils exhibit a dispersive movement without a clear tissue preference for relocation, consistent with previous reports [45, 79, 80]. This suggests that neutrophil reverse migration in zebrafish larvae does not appear to be influenced by the tissue where they initially migrated forward. Moreover, zebrafish neutrophils rarely undergo apoptosis at the injury sites [66, 79]. In our injury setup, we detected 1-2 % of neutrophils showing signs of irreversible death, and our RNA-seq data indicated ferroptosis as the mechanism responsible for this death. This is the first report of this type of spontaneous neutrophil death in zebrafish, previously observed in autoimmune disease models and the tumor microenvironment in mammals, where neutrophils become more immunosuppressive [81, 82]. Future studies on the implications of this mechanism of neutrophil cell death during tissue regeneration would be valuable for developing new tissue repair therapies.

Taken together, our observations challenge the classical view of neutrophils as “bad guys” and underscore their role as regulators of inflammation, whose functions need a full understanding to take advantage in specific biological situations. Moreover, our work also highlights the importance of choosing the right type of neutrophil-targeted therapy to enhance SCI regeneration and reduce potential side effects by demonstrating clear distinct phenotypes on SCI recovery after inducing decreased recruitment by Cxcr1/Cxcr2 inhibition and enhanced early resolution by Cxcr4 inhibition. Collectively, these results highlight the importance of the neutrophil inflammatory profile in spinal cord regeneration and suggest potential opportunities to modify the early inflammatory response to improve the regenerative capacity. Further investigation is needed to assess which neutrophil pro-resolution mediators are crucial to promote and enhance SCI regeneration and which mechanism trigger such phenotypes to develop better therapeutic approaches.

## ACKNOWLEDGEMENTS

We thank Yi Feng for sharing protocols, Anna Huttenlocher, Stephen A. Renshaw, Rita Fior and Anabela Bensimon-Brito for sharing zebrafish lines and Kimara L. Targoff for critical reading of the manuscript. We are grateful to the L. Saúde laboratory for constructive feedback. We thank A. Barros and L. Carvalho from the Zebrafish Unit, A. Nascimento, A. Temudo and J. Rino from the Bioimaging Unit and the Flow Cytometry Unit at the Instituto de Medicina Molecular João Lobo Antunes (iMM-JLA) for their expert advice and support. We are grateful to Adrien Jouary for sharing access to the zebrafish larvae behavioral set-up and for technical assistance. We thank Single Cell Discoveries for their bulk RNA-seq services and data analysis. This work was supported by a grant to C.d.S.-T. from the Fundação para a Ciencia e Tecnologia (FCT) (2022.02766.PTDC). C.d.S.-T. was supported by a contract from FCT (PTDC/BIA-BID/28572/2017). A.L and M.O were supported by grants from the Volkswagen Stiftung “Life?” Initiative and the ERC (Neurofish 773012). L.S. was supported by a FCT CEEC Individual Principal Investigator contract (2021.02253.CEECIND).

## AUTHOR CONTRIBUTIONS

C.d.S.-T. and L.S. conceived the project and provided funding; C.d.S.-T., L.R.L. and P.N.T. performed experiments; C.d.S.-T. and L.R.L. analyzed the data; C.d.S.-T. performed bioinformatic analysis and produced figures; A.L. and M.O. generated the tools to analyze larval swimming behavior; C.d.S.-T. wrote the initial draft; C.d.S.-T., L.R.L., S.d.O and L.S. reviewed and edited the manuscript.

---

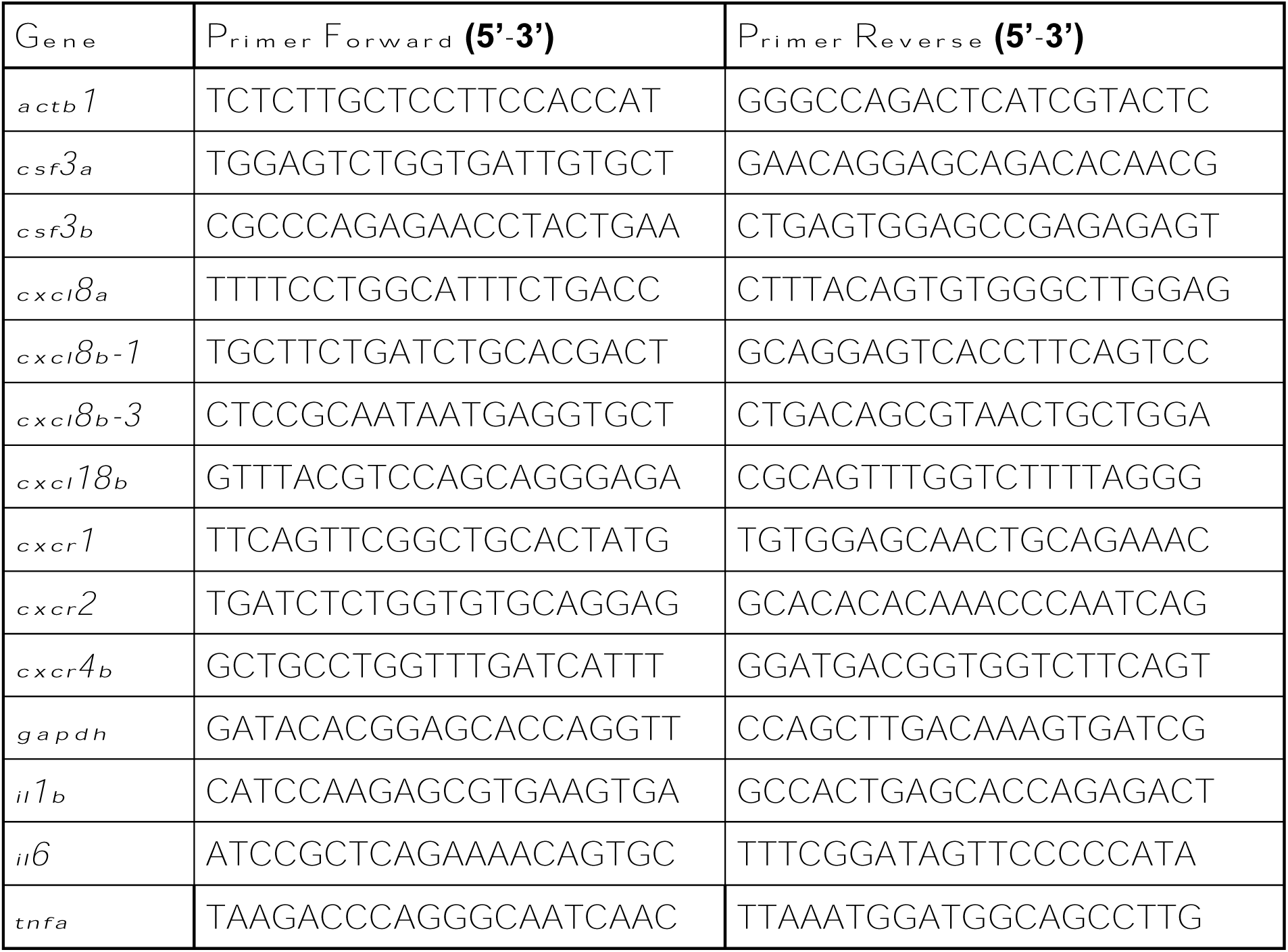
Supplementary Table 1 - qPCR primer sequences.

**Supplemental Movie 1: Representative movie showing neutrophils reverse migrating from the injury site throughout zebrafish body.**

**Figure S1:**
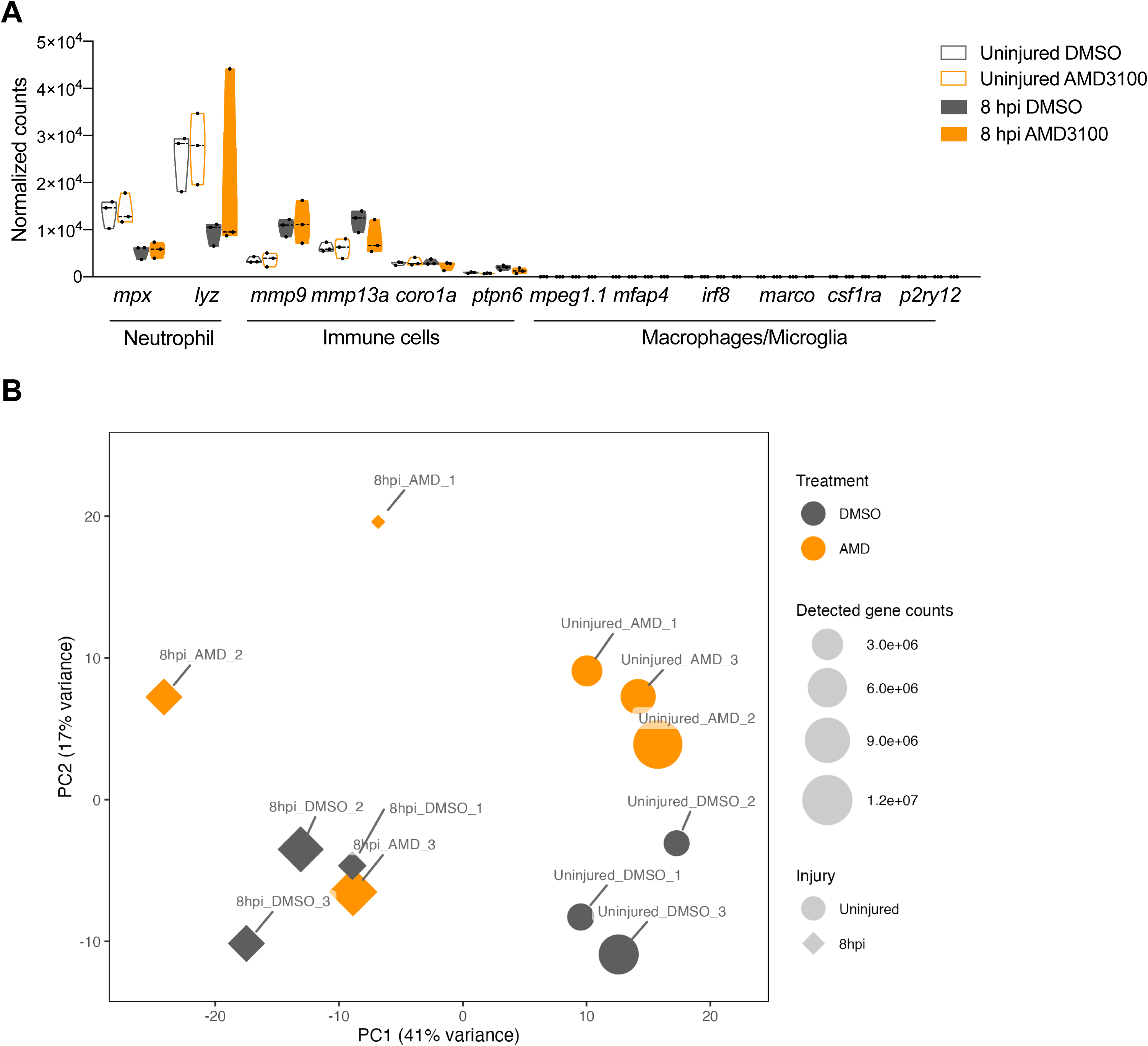
Principal component analysis of isolated neutrophils from AMD3100-treated larvae identifies injury (PC1) and treatment (PC2) variables. **(A)** Normalized RNA-seq counts of genes expressed by neutrophils, immune cells and macrophages/microglia in all four conditions. **(B)** PCA plot illustrates the clustering of RNA-seq samples with triplicates for each condition.

**Figure S2:**
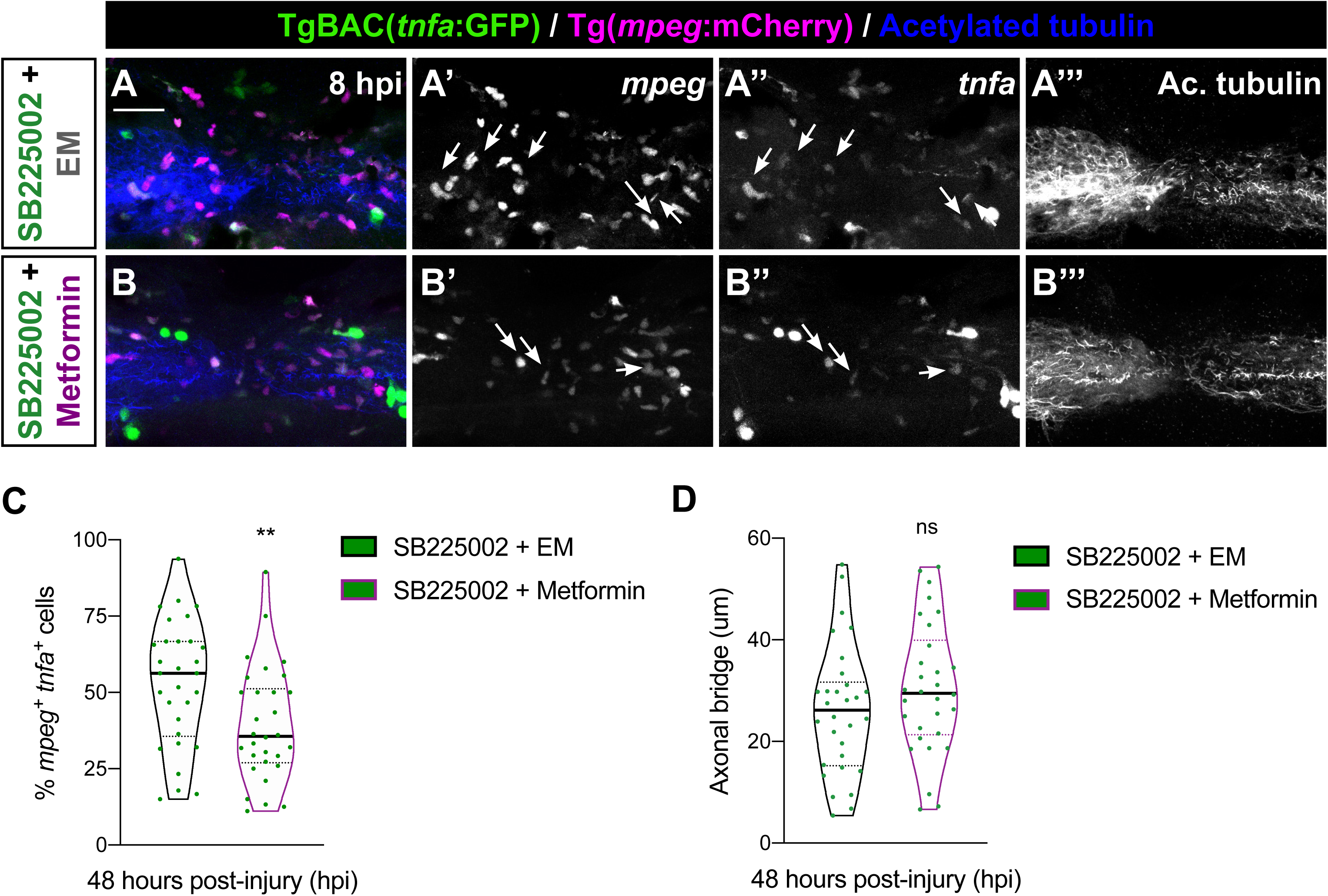
No improvement in axonal bridging in SB22502-treated larvae after Metformin treatment. **(A – B)** Representative maximal projections illustrate *mpeg^+^ tnfa^+^* cells in the injured spinal cord from SB225002-treated larvae followed by Metformin or embryo media incubation from 24 to 48 hpi. Individual panels for *tnfa* (A’, B’), *mpeg* (A’’, B’’) and acetylated tubulin (A’’, B’’’). Arrows highlight *mpeg^+^ tnfa^+^* cells. Scale bar, 30 μm. **(C)** Quantification of the proportion of *mpeg^+^ tnfa^+^* double positive cells over the total *mpeg^+^* population at 48 hpi (n = 30, conducted in four independent experiments). Unpaired two-tailed t-test demonstrates a statistically significant decrease in Metformin-treated group compared to the embryo media group (p =0.0095). **(D**) Quantification of the axonal bridge thickness revealed no statistically significant difference between Metformin- and embryo media-treated groups (p = 0.1992).

